# A Rarefaction Approach to Identify Local Introgression in a Three Population Tree

**DOI:** 10.64898/2026.05.13.724952

**Authors:** T. Quinn Smith, Zachary A. Szpiech

## Abstract

Patterson’s *D* statistic, also known as the *ABBA* − *BABA* statistic, is widely used to detect the presence of archaic genome-wide introgression between two non-sister taxa. Patterson’s *D* counts the imbalance between the number of biallelic sites where either the second and third taxa (ABBA site) share the derived allele or the first and third taxa (BABA site) share the derived allele in a four taxa tree. Here, the fourth taxon acts as an outgroup to determine the ancestral allele. When there is no introgression, these counts are expected to be equal, and a discordance between counts suggests introgression from the third taxon into either the first or second. Patterson’s D is limited to the detection of genome-wide introgression and exhibits a high false-positive rate when applied to smaller genomic segments. Here, we present a new method, D STatistic with Allelic Rarefaction (D*), to address these limitations. D* uses multiple lineages and does not require an outgroup to calculate the imbalance between the number of alleles found exclusively in the second and third taxa and the number of alleles found exclusively in the first and third taxa. D* employs a rarefaction technique to correct for unequal sample-size and allows multiallelic sites. We use simulations to show that D* has better precision and recall for detecting introgressed segments of DNA when compared to other methods. We conclude by recovering Denisovan DNA related to immune function in modern day Papuans.

## Introduction

Introgression is the transfer of genetic material between divergent lineages and is a pervasive demographic force that can shape the evolutionary trajectory of species. By introducing novel genetic variation, introgression can alter the genetic composition of populations, facilitate adaptation, and drive the speciation process Hibbins and Hahn (2022); Edelman and Mallet (2021). Recent genomic studies have highlighted the impact of introgression across human and non-human species, which has left detectable signatures in both modern and ancient populations Cahill et al. (2018); Edelman et al. (2019); Zhang et al. (2016); De Manuel et al. (2016); Green et al. (2010); Browning et al. (2018); Li et al. (2024).

Consequently, much effort has been dedicated to identifying and quantifying introgression. While classical population genetics models often struggle to capture the complexities of empirical data Whitlock and McCauley (1999), and likelihood methods are frequently too computationally intensive to generalize across large sample sizes or multiple populations Hey (2010); Gutenkunst et al. (2009); Kamm et al. (2020), summary statistics based on allele patterns across taxa are straightforward to compute and highly effective Patterson et al. (2012). A classic example of such a statistic is Patterson’s *D*, which can identify the presence of genome-wide introgression within a four-population tree by measuring imbalances in derived allele sharing Durand et al. (2011).

However, while Patterson’s *D* is robust when applied to genome-wide data in aggregate, the field’s focus has increasingly shifted toward locating specific, locally introgressed genomic segments Lopez Fang et al. (2024); Skov et al. (2018); Macià et al. (2025). In this context, Patterson’s *D* loses power, particularly in regions of low nucleotide diversity Martin et al. (2015). To address this, window-based adaptations such as *f*_*d*_ Martin et al. (2015) and *D*^+^ Lopez Fang et al. (2024), were developed.

Several other methods exist to locate introgressed segments in modern populations; however, such methods exploit linkage disequilibrium (LD) that is eroded over many generations, making them less reliable in detecting archaic segments Moorjani et al. (2011); Lawson et al. (2012). The statistics *S*\* Vernot and Akey (2014) and *SPrime* Browning et al. (2018) overcome the reliance on LD patterns but are model based and do not scale to many archaic samples. Similarly, likelihood-based methods Jacobs et al. (2019); Chen et al. (2020); Edelman and Mallet (2021) and approaches using the ancestral-recombination graph (ARG) Hubisz et al. (2020); Brandt et al. (2024); Nielsen et al. (2025) offer deep insights but sacrifice the computational simplicity of Patterson’s *D*, often scaling poorly with sample size. Other likelihood-based methods require phased genotypes Yuan et al. (2021), are unable to specify the archaic population, and require an estimate of mutation rate Macià et al. (2026).

The difficulty of identifying local introgressed segments has recently motivated the application of machine learning (ML) classifiers Ray et al. (2024); Cobb and Smith (2025); Durvasula and Sankararaman (2019, 2020). However, a major limitation of ML is its reliance on extensive simulated training data, which requires *a priori* knowledge of the demographic history of the populations in question Schrider and Kern (2018). Notably, these ML models consistently identify the number of private alleles within a genomic window as one of the most powerful predictive features for introgression Ray et al. (2024); Cobb and Smith (2025); Durvasula and Sankararaman (2020).

Private alleles are alleles found exclusively in one population and nowhere else. In the absence of gene flow, divergent populations accumulate private alleles independently; whereas, introgression increases allele sharing and reduces the number of private alleles. Slatkin first formalized the relationship between private alleles and migration rates Slatkin (1985); Barton and Slatkin (1986); Takahata and Slatkin (1986), although he noted that private allele counts are highly sensitive to sample size Barton and Slatkin (1986). Later, Kalinowski Kalinowski (2004) applied the rarefaction technique Hurlbert (1971) to correct for sample size bias. Rarefaction accomplishes this by calculating the probability of observing an allele in a population from a given sample size. The probability that an allele is private to a population is the probability that at least one copy of the allele exists in the population and does not exist in the other populations for a given sample size. The probabilities are summed over all alleles to give the sample size corrected number of private alleles. This approach was adapted to derive a generalization of private alleles to combinations of populations Szpiech et al. (2008).

Building on the theoretical foundations of Szpiech et al. (2008) Szpiech et al. (2008), we introduce the D STatistic with Allelic Rarefaction (*D*\*) which combines the reasoning behind *D/D*^+^ with generalized private alleles and rarefaction to identify the presence of introgression within local regions. While Patterson’s *D* relies on imbalances in derived allele counts at biallelic sites within a four-population tree Durand et al. (2011), an imbalance between counts indicates gene flow between populations within a pair. *D*\* generalizes this reasoning to sites with any number of alleles and removes the requirement of an outgroup. Indeed, recent work has demonstrated that outgroups are not strictly necessary for detecting genome-wide introgression Hahn and Hibbins (2019). Furthermore, by incorporating allelic rarefaction, *D*\* corrects for unequal sample sizes that might otherwise confound empirical studies.

In this study, we evaluate *D*\* through extensive simulations under different sample sizes, molecular clock violations, and unequal drift. We compare *D*\*’s performance to *D, D*^+^, *Sprime* Browning et al. (2018), and *hmmix* Macià et al. (2026) *in locating locally introgressed regions. For biallelic sites and a single lineage sampled from each population, D*\* performs identically to *D*^+^. Then, we show that *D*\*’s precision is greater than *D, D*^+^, *Sprime*, and *hmmix*’s precision when detecting locally introgressed regions. We show that neither *D*\*’s precision or recall when detecting locally introgressed regions or power to detect genome-wide introgression changes when sites are depolarized, unlike the other statistics. To ensure its utility for ancient DNA (aDNA) research, we test the performance of *D*\* against common empirical artifacts such as missing genotypes, deamination, and pseudohaploidization. Finally, we apply *D*\* to empirical data to identify previously known and novel Denisovan introgressed tracts in modern-day Papuans.

### Definition of Patterson’s *D* and *D*^+^

Consider a set of four taxa, {*P*_1_, *P*_2_, *P*_3_, *P*_4_} with a divergence relationship of (((*P*_1_, *P*_2_), *P*_3_), *P*_4_), indicating that *P*_1_ and *P*_2_ split after divergence from *P*_3_ and *P*_4_ is an outgroup. Considering only biallelic sites, let *A* denote the ancestral allele determined by *P*_4_ and let *B* denote the derived allele. At the *l*^*th*^ locus, we define the indicator variables *C*_*ABBA*_(*l*), *C*_*BABA*_(*l*), *C*_*BAAA*_(*l*), and *C*_*ABAA*_(*l*), to indicate the allele patterns (((*A, B*), *B*), *A*), (((*B, A*), *B*), *A*), (((*B, A*), *A*), *A*), and (((*A, B*), *A*), *A*) across the four taxa, respectively. Over *L* sites, Patterson’s *D* is defined as

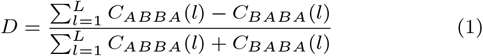

In the absence of introgression, *D* has an expected value of 0. If introgression occurred between *P*_2_ and *P*_3_, *D* is expected to be greater than 0. If introgression occurred between *P*_1_ and *P*_3_, *D* is expected to be less than 0 Durand et al. (2011). As the power of *D* to detect introgression in short segments, especially those with low nucleotide diversity, is weak Martin et al. (2015), *D*^+^ Lopez Fang et al. (2024) was developed to overcome this shortcoming. *D*^+^ is defined as

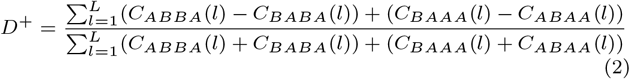

The interpretation of *D*^+^ is identical to *D*. If multiple lineages are available, let *p*_*li*_ be the derived allele frequency at the *l*^*th*^ site in the *i*^*th*^ population. Then, *D* and *D*^+^ are defined as

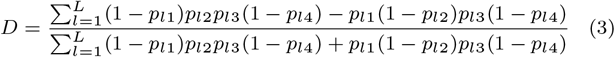

and

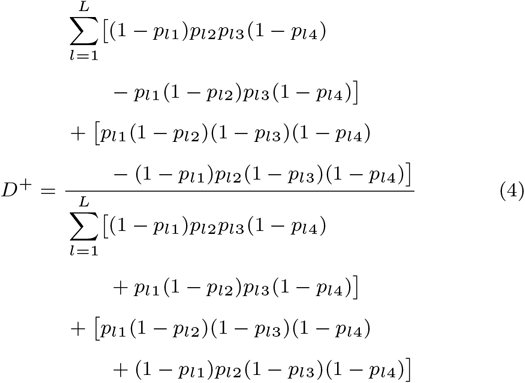

An important consideration is that the ancestral allele in Equation 1 and Equation 2 is determined by *P*_4_, while *p*_*li*_ are observed frequencies Durand et al. (2011); Malinsky et al. (2021). The derived allele frequencies from *P*_4_ are essential for controlling the false-negative rate of *D* and *D*^+^.

### Counting Alleles

We follow the approach of Szpiech et al. Szpiech et al. (2008) for counting alleles “private to combinations of populations”, that is, for counting alleles found in each of a set of populations and in no other population outside of that set. At a locus with *I* distinct alleles, we sample *N*_*ij*_ copies of allele *i* from population *j*, where 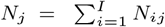 is the total number of lineages in population *j*.

We define the probability of *not* observing a copy of allele *i* from a subsample of size *g* lineages from population *j* as

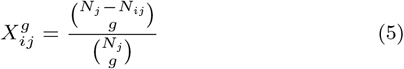

We calculate the probability of observing *at least* one copy of allele *i* from a subsample of size *g* lineages from population *j* as 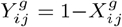. Then the number of alleles found in both *P*_1_ and *P*_3_ but absent in *P*_2_ in a subsample of size *g* lineages is given by

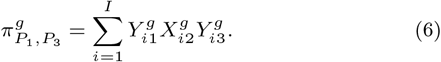

We call this term the number of alleles private to *P*_1_ and *P*_3_.

Similarly, the number of alleles common to *P*_2_ and *P*_3_ but absent in *P*_1_ in a subsample of size *g* lineages is given by

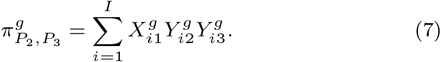

We call this term the number of alleles private to *P*_2_ and *P*_3_.

We also compute the total number of alleles at a locus (allelic richness) for populations *P*_1_, *P*_2_ and *P*_3_ combined. Let *N* = *N*_1_ + *N*_2_ + *N*_3_, then, for each allele, the probability of observing *at least* one copy of allele *i* in a sample size of *g* is 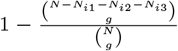. The total number of alleles at a locus is then given by

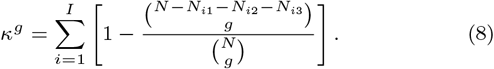

### Definition of *D*\*

Let 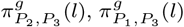, and *κ*^*g*^(*l*) be the statistics defined in Section 1.1 calculated at a locus *l*. We define *D*\* over *L* loci as

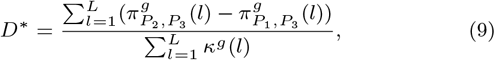

which contrasts the number of alleles private to *P*_1_ and *P*_3_ to the number of alleles private to *P*_2_ and *P*_3_, normalized by the allelic richness of the three populations combined. For biallelic sites and single lineages, we expect *D*\* (Equation 9) to be equivalent to *D*^+^ (Equation 2). Our reasoning is as follows. In Equation 9, 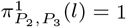 when *P*_2_ and *P*_3_ share an allele not found in *P*_1_. This corresponds to *C*_*ABBA*_(*l*) + *C*_*BAAA*_(*l*) in Equation 2. Likewise, 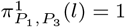 when *P*_1_ and *P*_3_ share an allele not found in *P*_2_, and this corresponds to *C*_*BABA*_(*l*)+*C*_*ABAA*_(*l*). Therefore, the numerators of *D*^+^ and *D*\* are the same. For the denominator of *D*\* in this case, *N* = 3, *N*_*i*1_ = 1, *N*_*i*2_ = 1, and *N*_*i*1_ = 1, resulting in *κ*^1^ = *C*_*BABA*_(*l*) + *C*_*ABAA*_(*l*) + *C*_*ABBA*_(*l*) + *C*_*BAAA*_(*l*) = 1. This shows that the denominators of Equation 2 and Equation 9 are the same, and therefore, shows that Equation 2 and Equation 9 are equivalent for the single lineages and biallelic loci.

In the case of no introgression from *P*_3_, *P*_1_ and *P*_2_ should share approximately equal amounts of alleles private with *P*_3_, that is, those inherited from the common ancestor of all three populations but lost in either *P*_1_ or *P*_2_ or those shared with *P*_3_ due to incomplete lineage sorting. In this case, we expect 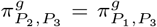 and *D*\* = 0. The rarefaction sample size correction maintains the expected result of *D*\* = 0 in the absence of introgression when multiple lineages are sampled from populations with the same effective population size. We provide empirical evidence of this in Table S1. In the case of gene flow between *P*_3_ and *P*_1_, we expect more alleles private to *P*_3_ and *P*_1_ relative to the number of alleles private to *P*_3_ and *P*_2_ and thus expect 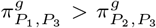 and *D*\* *<* 0. In the case of gene flow between *P*_3_ and *P*_2_, we expect more alleles private to *P*_3_ and *P*_2_ relative to the number private to *P*_3_ and *P*_1_, thus 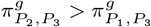 and *D*\* *>* 0.

We only calculate Equation 9 for sites where 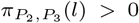 or 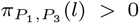, as including sites that are private to neither pair artificially deflates *D*\*. The parameter, *g*, is set to the minimum number of non-missing lineages within one of the three populations at a given site. *D*\* is calculated in non-overlapping genomic windows measured in base pairs.

### Comparing to *hmmix*

Given two sister populations *P*_1_ and *P*_2_, suppose *P*_1_ has negligible admixture from archaic populations and *P*_2_ is suspected of introgression with archaic populations. *hmmix* Macià et al. (2026) identifies segments in *P*_2_ introgressed from the archaic populations using a hidden Markov model (HMM) and variants private to *P*_2_ with respect to *P*_1_. The reason being that regions with a high concentration of variants in *P*_2_ not found in *P*_1_ are suspected to originate from the archaic populations. Then, the HMM is trained to classify non-overlapping segments as archaic or modern depending on the number of private alleles found within the segments.

*D*\* and *hmmix* differ in several ways. First, *hmmix* does not incorporate a sample size correction for the discovery of private alleles. Second, *hmmix* uses two populations and requires addition analyses to determine the source archaic population of an introgressed segment. Finally, each sample within *P*_2_ must be decoded individually using the trained HMM. To compare *hmmix* to the other methods, we trained the model using the samples from *P*_1_ and decoded each sample in *P*_2_. The decoding gave us the posterior probability a sample’s local segment is from the archaic population for each 1000 bp non-overlapping block along the simulated sequence. The posterior probabilities for each 1000 bp block were averaged across all samples, and then, averaged over fifty consecutive blocks. This gave us an average probability that non-overlapping 50 Kb blocks originated from the archaic population. We treated this value as a ‘statistic’ produced by *hmmix*.

### Comparing to *Sprime*

Another class of statistics used to identify local introgression from archaic populations are the *S*-statistics Li and Akey (2019). The original statistic being *S*\*Plagnol and Wall (2006), and the most recent being *Sprime* Browning et al. (2018), which generalized *S*\* to more samples, removed the requirement to window the genome, and is capable of identifying multiple waves of introgression. *Sprime* is similar to *hmmix* by requiring two populations: the first population must have little suspected archaic admixture and the second population is suspected of archaic admixture. *Sprime* employs a dynamic programming algorithm to identify ancestry found in the second population but absent in the first population. *SPrime* does not allow for the statistic to be calculated on defined windows. The *Sprime* program Browning et al. (2018) produces scores (the statistic’s value) of introgressed segments. A greater score indicates a greater belief that the segment was introgressed. Therefore, we averaged the score over all loci within 50 Kb, non-overlapping blocks to allow for comparison to *D, D*^+^, *D*\*, and *hmmix*.

### Performance Evaluation

We evaluated *D, D*^+^, *D*\*, *hmmix*, and *Sprime* in 50 Kb, non-overlapping blocks for all simulations. The block size of 50 Kb was chosen to reflect the average length of an archaically introgressed haplotype in our simplified demographic model Durvasula and Sankararaman (2019). If a statistic was undefined for any block, then the block was discarded. For each simulation scenario, we generated 100 replicates without gene flow (*f* = 0) to create null-distributions for the three statistics. The null-distributions were used to calibrate false-positive rates (FPRs). We treated 0 *< α <* 1 as our significance level. Since we are testing introgression between *P*_2_ and *P*_3_, we define our significance threshold at the top 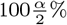 of values for the given statistic.

A true positive (TP) is a block that is statistically significant and introgressed from *P*_3_. In the case of sampling single lineages, we label a block as introgressed in the event that the lineage from *P*_2_ contains a total of 5 Kb from *P*_3_. In the case of sampling multiple lineages, we label a block as introgressed by satisfying two conditions. The first condition is that 10% of the sampled lineages must share a common segment from *P*_3_. The second condition is that the sum total of these shared segments must span at least 10% of the block (5 Kb). These requirements are identical to Fang et al Lopez Fang et al. (2024). A false negative (FN) is a block that was introgressed but not deemed statistically significant. A false positive (FP) is a block that was not introgressed but deemed statistically significant. We measure precision as 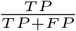, which is the probability that a block was introgressed and significant out of all significant blocks. We measure recall as 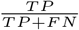, which is the probability that a block was introgressed and significant out of all introgressed blocks.

### Genome-Wide Introgression

Although the intent for *D*\* is to identify possibly introgressed local segments, it is worth investigating *D*\*’s ability to indicate genome-wide introgression. Our approach implements a block bootstrap to test the statistical significance of the genome-wide signal, unlike previous methods based on the rarefaction of private alleles Kalinowski (2004); Szpiech et al. (2008). *D*\* can accommodate multiple lineages sampled from the three populations in addition to incorporating multiallelic sites.

The significance of *D, D*^+^, and *D*\* was calculated using a block bootstrap with 1000 replicates Hahn and Hibbins (2019). For each statistic, the mean and standard deviation were calculated and used to parameterize a *z*-distribution to calculate significance. We set a significance threshold of *p <* 0.05 to determine the power of each statistic. The test is two-sided since it is unknown which pair of populations introgressed in practice. *hmmix* and *Sprime* are not intended to identify genome-wide introgression and excluded from genome-wide evaluation.

### Simplified Demographic Scenario

We used *msprime* Kelleher et al. (2016) to simulate a simplified demographic history (Figure 1) as a benchmark to model archaic introgression in modern humans, following previous work Racimo et al. (2017); Lopez Fang et al. (2024). We treated *P*_1_ and *P*_2_ as modern populations and *P*_3_ as the population that exchanges lineages with *P*_2_. We set the admixture proportion between *P*_3_ and *P*_2_ to *f* = 3% occurring at *T*_*GF*_ = 1, 600 generations ago. The divergence times between *P*_1_ and *P*_2_ is *T*_12_ = 4, 000 generations ago. The ancestral population, *P*_3_ diverged from *P*_1_ and *P*_2_ *T*_123_ = 16, 000 generations ago. We added an outgroup, *P*_4_, which diverged from the other three populations *T*_1234_ = 20, 000 generations ago. The effective population size for all populations was set to *N*_*e*_ = 10, 000. We set the length of the genomic region to 20 Mbp, the mutation rate to *µ* = 1.5×10^−8^ per base pair per generation, and the recombination rate to *ρ* = 1 × 10^−8^.

**Figure 1.**
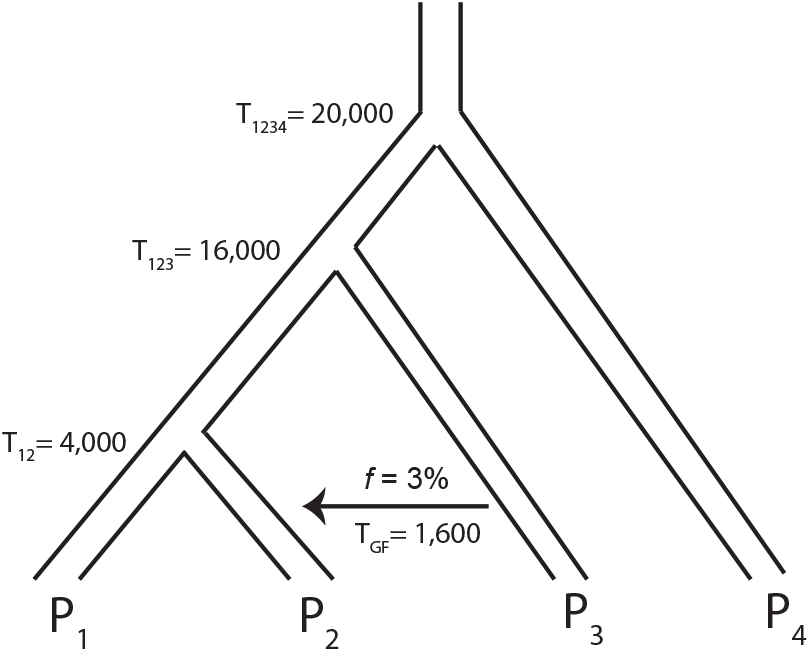
Simulated Demographic Scenario. The simplified demographic history used to evaluate *D*\*. Divergence times are in generations. By default, all populations have *N*_*e*_ = 10, 000.

First, we sampled a single lineage from each population and show the equivalence between *D*\* and *D*^+^. Neither *hmmix* or *Sprime* gave meaningful results with single lineages. We evaluated *D, D*\*, *D*^+^, *hmmix*, and *Sprime* sampling *N*_1_ = {1, 2, 6, 12} samples from *P*_1_, *N*_2_ = {1, 2, 6, 12} samples from *P*_2_, a single sample from *P*_3_, and a single sample from *P*_4_.Unless otherwise stated, all samples are assumed to be diploid.

### Depolarization

An advantage of *D*\* over *D* and *D*^+^ is that it does not require the outgroup population. The outgroup is used to determine the ancestral allele at each locus, a process known as polarization Voight et al. (2006). An ancestral allele at each site is required for Patterson’s *D* because it controls the false-negative rate of the statistic for genome-wide introgression detection Durand et al. (2011). In addition, the coalescent-based argument that supports the use of shared ancestral variation for identifying local regions of introgression relies on a known ancestral allele at each site Lopez Fang et al. (2024). However, a site’s ancestral allele is often unknown or requires a separate analysis to accurately determine its state. Studies often simplify the process and treat the reference allele as the ancestral allele Lopez Fang et al. (2024), possibly leading to a reduction in statistical power for both *D* and *D*^+^. In addition, choosing an outgroup too distant from the other populations may artificially inflate introgression detection Pang et al. (2025).

We tested the impact of removing polarization (depolarization) on both local introgression detection and genome-wide introgression detection for *D, D*^+^, and *D*\*. In our simulations, *P*_4_ acts as the outgroup population. We used EGGS Smith et al. (2026) to randomly swap the reference and alternative allele at each site with probability 0.5. This has the same effect as randomly changing the ancestral and derived alleles determined the outgroup (*P*_4_). Although unrealistic when an outgroup exists in the data, this approach removes all polarization information and allows us to investigate the sensitivity of the statistics to polarization error. We compared precision and recall for the detection of local introgression and power for genome-wide introgression.

### Molecular Clock Violations and Effects of Drift

An assumption of our simplified demographic model is the use of a constant mutation rate across all populations. Often, mutation rates vary between species, and therefore, affect the accumulation of derived alleles within a population Hodgkinson and Eyre-Walker (2011). This is in contrast to an identical mutation rate across all populations which influences the number of derived alleles evenly. Deviations in the molecular clock cause an imbalance in derived alleles between populations. For example, if one population accumulates derived alleles at a higher rate compared to other populations, then the number of private alleles in the first population will increase. This could result in a false signal of introgression Pang et al. (2025). *msprime* is not capable of varying mutation rates between populations directly. Instead, we mimic the uneven accumulation of mutations within a population by rescaling divergence times and sampling times Lopez Fang et al. (2024). Generally, if we want to scale the mutation rate of *P*_2_ by a factor *λ*, we increase all divergence times and *T*_*GF*_ by (*λ*−1)*T*_12_. Then, we sample *P*_2_ at time 0 and all other populations (*λ*−1)*T*_12_ generations ago. We can swap the roles of *P*_1_ and *P*_2_ to equivalently increase the mutation rate of *P*_1_ by a factor of *λ*. These scenarios were evaluated with a single sample from each population and considered *λ* = {1.2, 1.5, 2, 5} for *P*_1_ and *P*_2_ Pang et al. (2025).

Genetic drift is the random change in allele frequencies over time as a consequence of finite population size Gillespie (2004). In small populations this leads to a higher rate of fixation or loss compared to large populations. We expect unequal drift between *P*_1_ and *P*_2_ to influence *D*\* since a large discordance in drift could cause unequal accumulation of private alleles. We evaluated the effects of drift by varying *N*_*e*_ = {2500, 5000, 25000} for *P*_1_ and *P*_2_, creating six different scenarios. We kept the other populations’ effective population size at *N*_*e*_ = 10000. These scenarios were evaluated with a single sample from each population.

### Missing Data, Deamination, Pseudohaploidization

Simulated data do not contain sources of error known to bias inference Sethuraman (2026); Dabney et al. (2013). We also simulate several sources of error on top of our simulated genomic region that are common when working with modern and ancient DNA. We used parameters gathered from ancient DNA studies to maintain a sense of realism and act as a baseline for other studies.

The low quality of aDNA prevents many alleles from being accurately called, which leads to a large amount of missing data. When an allele can be called, the coverage is often too low to confidently call a second allele Sousa da Mota et al. (2023). For aDNA samples that are possibly heterozygous at a site, a random allele is chosen from the reads, and the sample is forced to be homozygous for the random allele. This process is known as pseudohaploidization Jakobsson and Mattias (2019). In addition, aDNA is particularly prone to the deamination of cysteine to thymine Dabney et al. (2013); Harney et al. (2021). We investigated the influence of all three processes on *D*\*.

We used the simplified demographic history and sampled 12 individuals from *P*_1_, 12 individuals from *P*_2_, 2 individuals from *P*_3_, and 1 individual from *P*_4_. A sample’s genotype is missing at a site if both alleles are absent (./.). We introduced missing genotypes according to the process described by Pandey et al Pandey et al. (2024). Define a *β*-distribution with a mean of 0.55 and a standard deviation of 0.23. For each sample’s genotype, we drew a random number, *b*, from our *β*-distribution to determine the proportion of individuals that will be treated as missing at the given site. We randomly set both alleles missing (./.) for ⌊27*b*⌋ of the individuals. We introduced deamination according to Harney et al Harney et al. (2021) and treated a locus as a transition with a probability of 77.6%. For each alternative allele, we switched it to the reference allele with a probability of 5%. Then, pseudohaploids were created in the dataset as follows. For each heterozygous individual, we randomly selected one of the alleles with equal probability to make the individual homozygous for that allele Jakobsson and Mattias (2019). We investigated the effects of all three mechanisms separately and together. Missing genotypes, deamination, and pseudohaploization were introduced into the replicates using the EGGS software Smith et al. (2026). We limited to the analysis to *D, D*\*, and *D*^+^ because *Sprime* does not accommodate missing genotypes and *hmmix* requires tracks of missing genotypes to be masked out Macià et al. (2026).

### Denisovan Introgression in Papuans

There is extensive evidence of introgression from Denisovans into modern-day Papuan individuals Meyer et al. (2012); Malaspinas et al. (2016); Jacobs et al. (2019); Vespasiani et al. (2022). We used *D*\* to explore possibly introgressed Denisovan segments in Papuan individuals. Modern-day Sardinians possess low amounts of admixture with Papuans and are more recently diverged from Papuans than Denisovans Malaspinas et al. (2016). This results in the tree ((*Sardinians, Papuans*), *Denisovans*). Here, Sardinians, Papuans, and Denisovans act as *P*_1_, *P*_2_, and *P*_3_, respectively.

We retrieved the Denisovan sample from Meyer et al Meyer et al. (2012) that was later reprocessed in Prufer et al Prüfer et al. (2017) from v62 of the Allen Ancient DNA Resource Mallick et al. (2024). We converted the dataset to Variant Call Format using EGGS Smith et al. (2026). The sample’s variants were lifted over from hg19 to hg38. Then, we extracted the 27 Sardinian individuals and the 17 Papuan individuals from the whole-genome sequences of the Human Genome Diversity Project (HGDP) Koenig et al. (2024). These samples were merged with the Denisovan sample using bcftools Danecek et al. (2021). Only biallelic autosomal variants were included in the analysis. We excluded *Sprime* from the analysis since it does not accommodate missing genotypes.

### Performance under Simplified Demographic Scenario

In the case of biallelic sites and single haploids from each population, the ancestral allele determined by the outgroup population is used directly in the calculation of *D* and *D*^+^. Simulating single haploids under our simplified demographic scenario (Figure 1), we calculated precision and recall for *D, D*^+^, and *D*\* (Figure 2). The non-continuous appearance of *D*’s precision and recall curves is caused by a multi-modal, discrete null-distribution (Figure S1). The precision and recall for *D*^+^ and *D*\* are the same and outperform *D*. This supports our reasoning that *D*\* is equivalent to *D*^+^ in the case of biallelic sites and single lineages.

**Figure 2.**
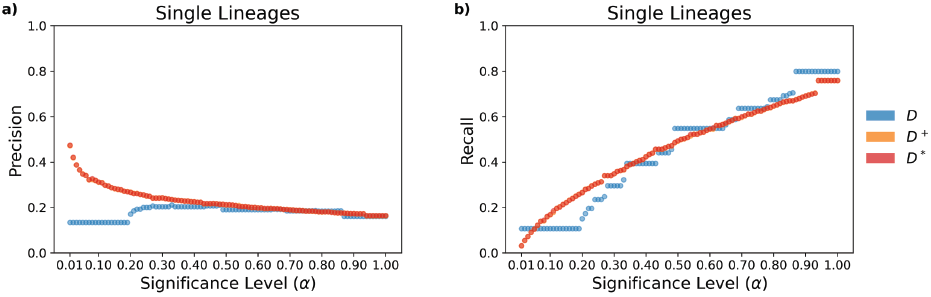
Results for single haploid lineages. **a)** Precision and **b)** recall are shown for significance levels (*α*) between 0.01 and 1.

When there are multiple lineages from the populations, *D* and *D*^+^ use derived allele frequencies (Equation 3 and Equation 4) instead of an explicitly labeled ancestral allele (Equation 1 and Equation 2), whereas *D*\* does not change its calculation when sample size grows beyond a single lineage per population. The rarefaction technique in *D*\* is expected to be most beneficial when samples are small and uneven. We therefore evaluated *D, D*^+^, *D*\*, *Sprime*, and *hmmix* on diploid samples of {1, 2, 6, 12} taken from *P*_1_ and *P*_2_ simulated under the simplified demographic scenario (Figure 1). We tested each combination sample sizes from *P*_1_ and *P*_2_ and a single diploid sample from *P*_3_ and *P*_4_, totaling 16 sampling configurations. We further evaluated these statistics when ancestral allele state information was removed. The results for a single diploid sample from *P*_1_ and *P*_2_ is shown in Figure 3, and the other combinations are shown in Figures S2-S16.

**Figure 3.**
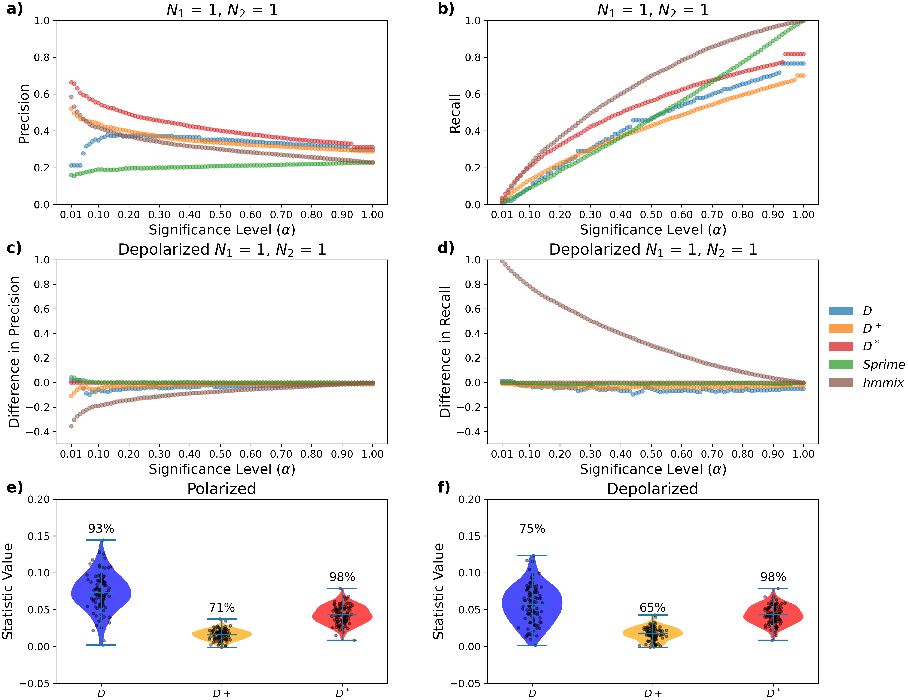
Results for diploid sampling. *P*_1_ = 1 **and** *P*_2_ = 1 **a)** Precision and **b)** recall are shown for significance levels (*α*) between 0.01 and 1. **c)** Difference (polarized minus depolarized) in precision and **d)** in recall. **e)** Genome-wide power for introgression detection on polarized genotypes, distribution gives local window values, percentage above gives power. **f)** Genome-wide power for introgression detection on unpolarized genotypes, distribution genome-wide values, percentage above gives power.

Figure 3a plots precision as a function of significance level and shows *D*\* outperforming the other statistics. Figure 3b plots recall as a function of significance level and shows *D*\* outperforming or tying the other statistics. In Table 1 we collect precision and recall for all statistics fixing the significance level at *α* = 0.05 for all 16 sample-size combinations. We see that *D*\* has better precision than the other statistics in 14 of the 16 combinations. In addition, *D*\* has better recall for all cases with a single sample from *P*_2_ and all cases with 12 samples from *P*_1_. *D*\* has better precision and recall than *D, D*^+^, and *Sprime* in all combinations. *hmmix* is the only statistic to challenge *D*\*. However, when the alleles are depolarized, we see that precision (Figure 3c) and recall (Figure 3d) for *hmmix* is the statistic with the largest change. The slight advantage of *hmmix* with polarized sites becomes a major disadvantage for unpolarized sites. We note that although recall for *hmmix* using depolarized alleles goes up substantially, precision drops similarly. This implies that although most of the truly introgressed segments are being called correctly, the false positive rate is also extremely high. *D*\* is the only statistic of the five that does not exhibit any change in precision or recall between polarized and unpolarized sites.

**Table 1.**
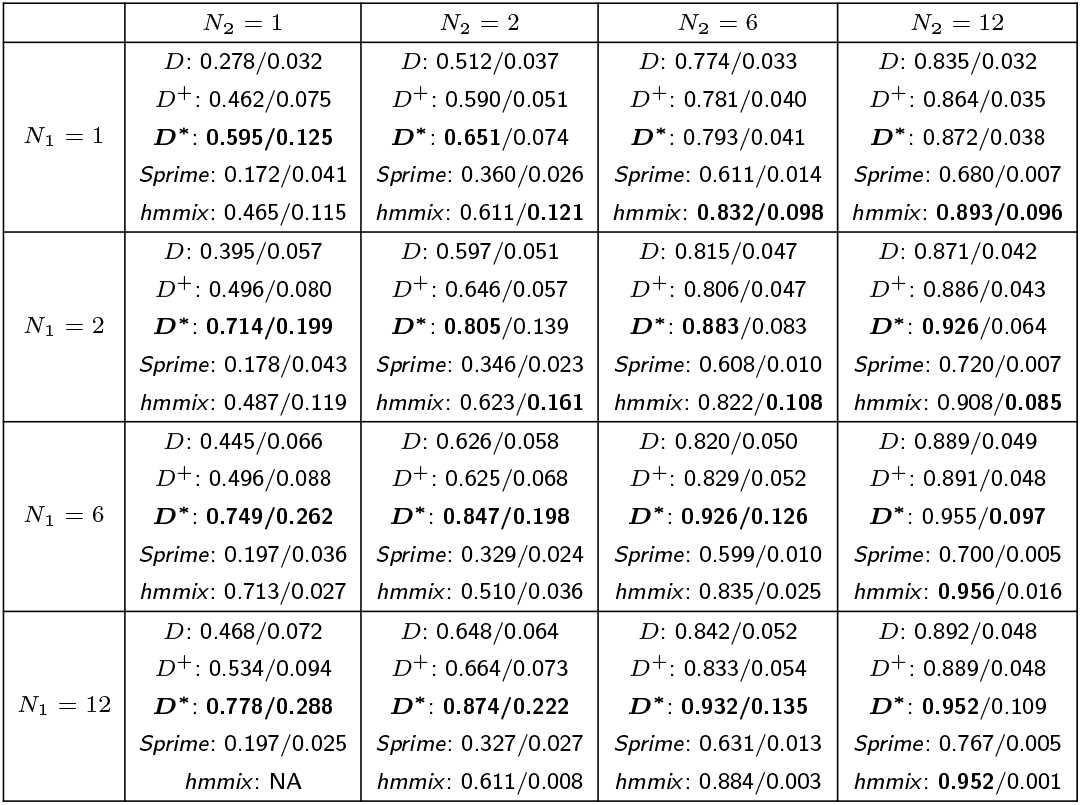
Precision and Recall for Different Sample Sizes at. *α* = 0.05 Precision and recall for identifying locally introgressed blocks for each combination of *P*_1_ = {1, 2, 6, 12} and *P*_2_ = {1, 2, 6, 12} at the significance level *α* = 0.05. The largest precision and recall for each combination among the statistics are printed in bold. Note, *hmmix* is undefined when *P*_1_ = 12 and *P*_2_ = 1.

|  | *N*<sub>2</sub> = 1 | *N*<sub>2</sub> = 2 | *N*<sub>2</sub> = 6 | *N*<sub>2</sub> = 12 |
|---|---|---|---|---|
| *N*<sub>1</sub> = 1 | *D*: 0.278/0.032<br>*D*<sup>+</sup>: 0.462/0.075<br>***D\****: **0.595/0.125**<br>*Sprime*: 0.172/0.041<br>*hmmix*: 0.465/0.115 | *D*: 0.512/0.037<br>*D*<sup>+</sup>: 0.590/0.051<br>***D\****: **0.651**/0.074<br>*Sprime*: 0.360/0.026<br>*hmmix*: 0.611/**0.121** | *D*: 0.774/0.033<br>*D*<sup>+</sup>: 0.781/0.040<br>***D\****: 0.793/0.041<br>*Sprime*: 0.611/0.014<br>*hmmix*: **0.832/0.098** | *D*: 0.835/0.032<br>*D*<sup>+</sup>: 0.864/0.035<br>***D\****: 0.872/0.038<br>*Sprime*: 0.680/0.007<br>*hmmix*: **0.893/0.096** |
| *N*<sub>1</sub> = 2 | *D*: 0.395/0.057<br>*D*<sup>+</sup>: 0.496/0.080<br>***D\****: **0.714/0.199**<br>*Sprime*: 0.178/0.043<br>*hmmix*: 0.487/0.119 | *D*: 0.597/0.051<br>*D*<sup>+</sup>: 0.646/0.057<br>***D\****: **0.805**/0.139<br>*Sprime*: 0.346/0.023<br>*hmmix*: 0.623/**0.161** | *D*: 0.815/0.047<br>*D*<sup>+</sup>: 0.806/0.047<br>***D\****: **0.883**/0.083<br>*Sprime*: 0.608/0.010<br>*hmmix*: 0.822/**0.108** | *D*: 0.871/0.042<br>*D*<sup>+</sup>: 0.886/0.043<br>***D\****: **0.926**/0.064<br>*Sprime*: 0.720/0.007<br>*hmmix*: 0.908/**0.085** |
| *N*<sub>1</sub> = 6 | *D*: 0.445/0.066<br>*D*<sup>+</sup>: 0.496/0.088<br>***D\****: **0.749/0.262**<br>*Sprime*: 0.197/0.036<br>*hmmix*: 0.713/0.027 | *D*: 0.626/0.058<br>*D*<sup>+</sup>: 0.625/0.068<br>***D\****: **0.847/0.198**<br>*Sprime*: 0.329/0.024<br>*hmmix*: 0.510/0.036 | *D*: 0.820/0.050<br>*D*<sup>+</sup>: 0.829/0.052<br>***D\****: **0.926/0.126**<br>*Sprime*: 0.599/0.010<br>*hmmix*: 0.835/0.025 | *D*: 0.889/0.049<br>*D*<sup>+</sup>: 0.891/0.048<br>***D\****: 0.955/**0.097**<br>*Sprime*: 0.700/0.005<br>*hmmix*: **0.956**/0.016 |
| *N*<sub>1</sub> = 12 | *D*: 0.468/0.072<br>*D*<sup>+</sup>: 0.534/0.094<br>***D\****: **0.778/0.288**<br>*Sprime*: 0.197/0.025<br>*hmmix*: NA | *D*: 0.648/0.064<br>*D*<sup>+</sup>: 0.664/0.073<br>***D\****: **0.874/0.222**<br>*Sprime*: 0.327/0.027<br>*hmmix*: 0.611/0.008 | *D*: 0.842/0.052<br>*D*<sup>+</sup>: 0.833/0.054<br>***D\****: **0.932/0.135**<br>*Sprime*: 0.631/0.013<br>*hmmix*: 0.884/0.003 | *D*: 0.892/0.048<br>*D*<sup>+</sup>: 0.889/0.048<br>***D\****: **0.952**/0.109<br>*Sprime*: 0.767/0.005<br>*hmmix*: **0.952**/0.001 |

We evaluate genome-wide introgression detection as well, although *Sprime* and *hmmix* are not intended for this task and are excluded from the analysis. We calculated genome-wide power for each population size combination for *D, D*^+^, and *D*\* (Figure 3e). On average, we found that *D*\* had similar to slightly better power when compared to *D* and higher power when compared to *D*^+^. The power for both *D* and *D*\* dropped on the depolarized set while the power for *D*\* remained unchanged (Figure 3f). We show the change in genome-wide power for every sample size combination in Table S2. Results across the other combinations of sample sizes (Figures S2-S16) were generally qualitatively similar.

### Performance under Molecular Clock Violations

We evaluated *D, D*^+^, *D*\*, *Sprime*, and *hmmix* under molecular clock violations by varying the mutation rate for each population, *P*_1_ and *P*_2_, by a factor of *λ* = {1.2, 1.5, 2, 5} with respect to the other population. Indeed, the mean for the null-distribution of *D*^+^ and *D*\* is non-zero for all molecular clock violations, but *D*\* maintains better precision in all cases. The precision and recall curves are shown in Figure S17 and Figure S18, respectively. The precision and recall at the significance level *α* = 0.05 for the five statistics is shown in Table 2. In each scenario, *D*\* exhibits a higher precision than the other statistics. *Sprime* or *hmmix* exhibit a better recall compared to *D*\*, except for the case of *P*_2_’s *λ* = 5. We note that even though violations of the molecular clock create a null *D*\* distribution with non-zero mean, the inference of local introgression remains powerful since we are taking an outlier approach with respect to the rest of the genome.

**Table 2.**
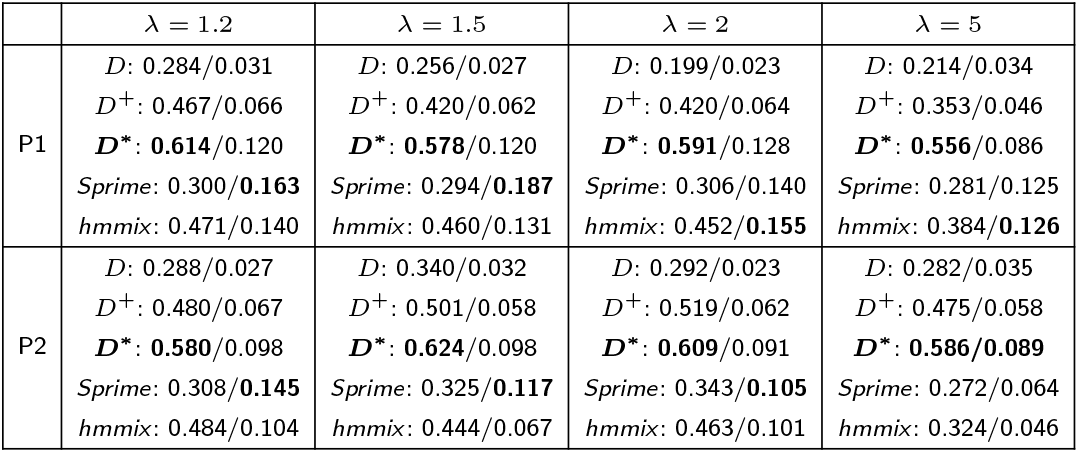
Precision and Recall for Molecular Clock Violations at. *α* = 0.05 Precision and recall for identifying locally introgressed blocks for scaling the mutation rate of *P*_1_ and *P*_2_ by a factor of *λ* = {1.2, 1.5, 2, 5} at the significance level *α* = 0.05. The largest precision and recall for each combination among the statistics are printed in bold.

|  | λ = 1.2 | λ = 1.5 | λ = 2 | λ = 5 |
|---|---|---|---|---|
| P1 | *D*: 0.284/0.031<br>*D*<sup>+</sup>: 0.467/0.066<br>***D<sup>\*</sup>***: **0.614**/0.120<br>*Sprime*: 0.300/**0.163**<br>*hmmix*: 0.471/0.140 | *D*: 0.256/0.027<br>*D*<sup>+</sup>: 0.420/0.062<br>***D<sup>\*</sup>***: **0.578**/0.120<br>*Sprime*: 0.294/**0.187**<br>*hmmix*: 0.460/0.131 | *D*: 0.199/0.023<br>*D*<sup>+</sup>: 0.420/0.064<br>***D<sup>\*</sup>***: **0.591**/0.128<br>*Sprime*: 0.306/0.140<br>*hmmix*: 0.452/**0.155** | *D*: 0.214/0.034<br>*D*<sup>+</sup>: 0.353/0.046<br>***D<sup>\*</sup>***: **0.556**/0.086<br>*Sprime*: 0.281/0.125<br>*hmmix*: 0.384/**0.126** |
| P2 | *D*: 0.288/0.027<br>*D*<sup>+</sup>: 0.480/0.067<br>***D<sup>\*</sup>***: **0.580**/0.098<br>*Sprime*: 0.308/**0.145**<br>*hmmix*: 0.484/0.104 | *D*: 0.340/0.032<br>*D*<sup>+</sup>: 0.501/0.058<br>***D<sup>\*</sup>***: **0.624**/0.098<br>*Sprime*: 0.325/**0.117**<br>*hmmix*: 0.444/0.067 | *D*: 0.292/0.023<br>*D*<sup>+</sup>: 0.519/0.062<br>***D<sup>\*</sup>***: **0.609**/0.091<br>*Sprime*: 0.343/**0.105**<br>*hmmix*: 0.463/0.101 | *D*: 0.282/0.035<br>*D*<sup>+</sup>: 0.475/0.058<br>***D<sup>\*</sup>***: **0.586/0.089**<br>*Sprime*: 0.272/0.064<br>*hmmix*: 0.324/0.046 |

We show the genome-wide power for *D, D*^+^, and *D*\* under molecular clock violations in Figure S19. Regardless of *P*_1_’s *λ, D*\* maintains equal or better power than *D* and *D*^+^ with the power of *D* decreasing as *λ* increases. This performance does not transfer to the case of changing *P*_2_’s *λ*. Both *D*^+^ and *D*\* suffer from accelerated mutation on *P*_2_’s lineage with power dropping to 0 for *λ* ≥ 1.5. *D* maintains power (*>* 90%) for all values of *λ*. This is unsurprising since a disproportionate increase in the mutation rate along *P*_2_ will cause a disproportionate accumulation of allele exclusive to *P*_2_. Therefore, the proportion of private alleles to the combination of *P*_2_ and *P*_3_ will decrease.

### Performance under Varying Drift

Next, we evaluated the statistics under varying levels of drift in *P*_1_ and *P*_2_ by changing either population’s *N*_*e*_ = {2500, 5000, 25000}. Similar to violations in the molecular clock, the mean of the null-distribution for *D*\* is non-zero (Table S4). However, for a significance level of *α* = 0.05, *D*\* maintains better precision than the other statistics for each scenario (Table 3). We show the precision and recall curves in Figure S20 and Figure S21, respectively. As with violations of the molecular clock, we note that even though varying drift regimes create a null *D*\* distribution with non-zero mean, the inference of local introgression remains powerful since we are taking an outlier approach with respect to the rest of the genome.

**Table 3.**
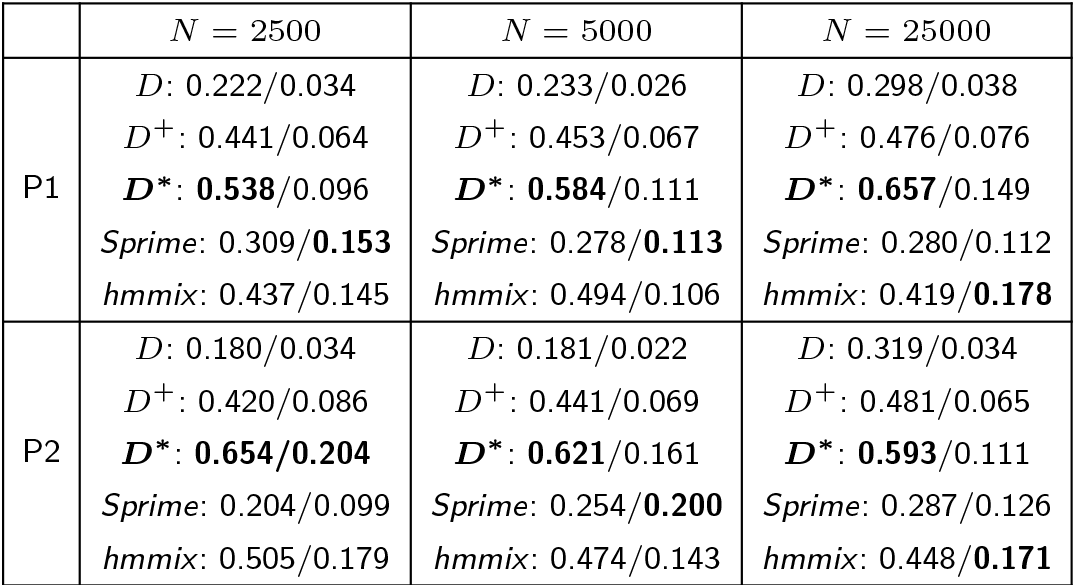
Precision and Recall for Varying Drift at. *α* = 0.05 Precision and recall for identifying locally introgressed blocks for varying drift in *P*_1_ and *P*_2_ by setting *N*_*e*_ = {2500, 5000, 25000} at the significance level *α* = 0.05. The largest precision and recall for each combination among the statistics are printed in bold.

We evaluated genome-wide power for *D, D*^+^, and *D*\* under the different drift scenarios (Figure S22). We found that when *P*_2_’s *N*_*e*_ was less than *P*_1_’s *N*_*e*_ (*N*_*e*_ = {2500, 5000}), the genome-wide power for *D*\* is 0 and *D* and *D*^+^ remain unaffected. *D*\*’s poor performance can be explained by alleles being lost or fixed due to the stronger drift in *P*_2_ compared to *P*_1_. When *P*_1_’s *N*_*e*_ = 25000, drift is weak and contains more diversity compared to *P*_2_. *D*\*’s power drops to 0 in this case because *P*_1_ is more likely to share alleles with *P*_3_ than the number of alleles shared between *P*_2_ and *P*_3_.

An important point to make is that *D* and *D*^+^ use derived allele frequencies in their calculations and can be interpreted as measuring discordance in drift between pairs of populations. This can explain the advantage of *D* and *D*^+^ compared to *D*\* for detecting genome-wide introgression in the presence of strong differences in drift. *D*\* uses allele frequencies implicitly through the rarefaction technique and is control by the standardized sample size *g*. Essentially, lowering *g* down-weights alleles at low frequencies in the calculation of *D*\*. For these computations, *g* = 2 at each site. We reran the genome-wide introgression detection for varying amounts of drift using *g* = 1, which shifts weight to common alleles. The results are plotted in Figure S23 showing *D*\*’s power is less than that of *D* but greater than *D*^+^’s power. Setting *g* = 1 shows that the genome-wide mean for *D*\* is 0 in the absence of introgression even in the presence of strong drift (Table S5). This suggests that the standardized sample size could be tuned to correct *D*\* for genome-wide introgression detection.

### Performance under aDNA Artifacts

Simulations lack many of the errors found in aDNA. So we simulated some of the typical artifacts observed when handling empirical aDNA in order to evaluate how these statistics perform. We simulated 10 diploid samples from *P*_1_, 10 diploid samples from *P*_2_, 2 diploid samples from *P*_3_, and 2 diploid samples from *P*_4_. *D, D*^+^, and *D*\* were evaluated since they are able to accommodate missing genotypes. We synthetically made all samples pseduohaploid and introduced synthetic deamination and missing genotypes. We recognize that pseudohaploidization of all samples effectively cuts the number of unique lineages in half. We include it for completeness since it is common among ancient samples. Typically, *D* and *D*^+^ are used to test introgression between modern-day populations and an archaic population, where the archaic population’s genotypes are pseudohaploid and contain deamination and missing genotypes. We introduced these artifacts across all samples to maintain homogeneity. We inspected the effects of pseudohaploids, demaination, and missing genotypes separately and together (Figure S24). *D*\* maintains better precision for *α <* 0.05 for all scenarios. *D*\*’s recall is better than that of *D* and *D*^+^ for pseudohaploids and denomination. Recall for all three statistics suffer due to missing genotypes, and *D* has slightly better recall than *D*^+^ and *D*\* when missing genotypes are included. Out of the three artifacts, missing genotypes has the largest impact on performance.

### Denisovan Introgression in Papuans

When a demographic history relating populations is known, then a null-distribution for *D*\* can be generated and *p*-values can be calculated for empirical data. When a demographic history is unknown, *D*\* is fundamentally an outlier test. Regions with a large *D*\*, in magnitude, compared to the genome-wide distribution could indicate that the region is introgressed. Therefore, *D*\* and *D*^+^ are exploratory statistics used to identify regions of interest for future analysis. We conducted an analysis on 27 Sardinian individuals, 17 Papuan individuals, and one pseudohaploid Denisovan assuming an unknown demographic history to illustrate an exploratory analysis on empirical data. We calculated *hmmix* on the dataset but excluded *Sprime* since it does not accomodate missing genotypes.

Of the top 0.1% of blocks with at least 10 sites for *D*^+^ and *D*\*, 9 were common to both sets (Figure 4a). We do not show the comparison to *D* and *hmmix* since no blocks were shared with either *D*^+^ or *D*\*. Table S6 shows the top 0.1% of blocks with at least 10 sites and the blocks’ overlapping protein coding genes unique to *D*\*. Table S7 and Table S8 show the top 0.1% of blocks with at least 10 sites and the blocks’ overlapping protein coding genes or *D*^+^ and *hmmix*, respectively. For *D*\*, *RRM* 1 Zhou et al. (2022), *HIC*2 Huang et al. (2022), *BANK*1 Gómez Hernández et al. (2021), *CLEC*9*A* Zhang et al. (2012), and *LRBA* Lopez-Herrera et al. (2012) are all related to immune function. This is consistent with expectations that genes related to the immune system are likely to be specifically introgressed from Denisovans into Papuans Vespasiani et al. (2022). We found *ADK* (Adenosine Kinase), which is a potential candidate for positive selection in Papuans and thought to originate from archaic ancestors González-Buenfil et al. (2024). *ADK* is also related to immune function Peter-Okaka and Boison (2025).

**Figure 4.**
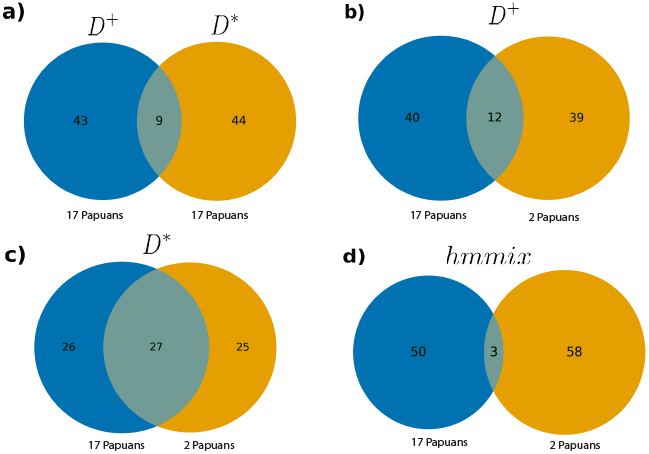
Overlap for Top 0.1% of Blocks. **a)** Blocks common and exclusive to *D*^+^ (blue) and *D*\* (orange) in the top 0.1% using all Papuans. **b)** Blocks common and exclusive to *D*^+^ in the top 0.1% using all Papuans (blue) and only 2 Papuans (orange). **c)** Blocks common and exclusive to *D*\* in the top 0.1% using all Papuans (blue) and only 2 Papuans (orange). **d)** Blocks common and exclusive to *hmmix* in the top 0.1% using all Papuans (blue) and only 2 Papuans (orange).

On the simulated data, *D*\* performed well for unequal sample sizes, suggesting that the rarefaction approach provides an advantage. We tested this on the empirical data. We reran the analysis using 27 Sardinian individuals, two Papuan individuals, and the pseudohaploid Denisovan. We compared the top 0.1% of blocks for this down-sampled set to the set found using all 17 Papuan individuals. *D*^+^ shared 12 blocks between the full and down-sampled sets (Figure 4b), *D*\* shared 27 blocks between the full and down-sampled sets (Figure 4c), *hmmix* shared 3 blocks between the full and down-sampled sets (Figure 4d). For *D*\*, the blocks overlapping *RRM* 1 and *LRBA* were retained after down-sampling. Since *D*\* retains more blocks compared to *D*^+^ and *hmmix* when down-sampling, we have evidence that the sample size correction employed by *D*\* is advantageous in empirical data.

## Conclusion

Patterson’s *D* statistic is widely used to detect the presence of genome-wide archaic admixture Durand et al. (2011). Recent work has focused on identifying introgressed regions from archaic populations in modern-day populations Martin et al. (2015); Skov et al. (2018); Lopez Fang et al. (2024). In a three population tree with an unadmixed ancestral outgroup, Patterson’s *D* is calculated from the difference between the number of sites where the first and third population share the derived allele and the second and third population share the derived allele. An imbalance of counts indicates the presence of genome-wide introgression. *D* has a high false-positive rate in regions of low nucleotide diversity Martin et al. (2015). *D*^+^ addressed this by incorporating sites where either pair of populations share the ancestral allele. When populations have multiple lineages, the counts are replaced with derived allele frequencies.

Here, we presented a new approach combining the simplicity of Patterson’s *D* Patterson et al. (2012) and the information captured by a rarefaction technique to count alleles private to pairs of populations Szpiech et al. (2008). We evaluated *D, D*^+^ Lopez Fang et al. (2024), *Sprime* Browning et al. (2018), *hmmix* Macià et al. (2026), and *D*\* using simulations generated under a simplified demographic model resembling archaic introgression (Figure 1). In the case of biallelic sites and single lineages, *D*\* is equivalent to *D*^+^. For multiple samples, *D*\* had better precision than the other statistics in most circumstances, including molecular clock violations and varying amounts of drift. In addition, we showed that the precision and recall for *D*\* did not change for depolarized genotypes unlike for *D, D*^+^, *Sprime*, and *hmmix*. We examined *D*\* in the presence of errors common to aDNA, such as pseudohaploids, deamination, and missing genotypes, and showed that while *D*\*’s performance did suffer, *D*\*’s inference is still reliable compared to other methods for local introgression detection.

Patterson’s *D* requires an outgroup to establish the ancestral allele at each site, a process known as polarization, to control the false-negative rate for genome-wide introgression detection. Removing the need for polarized alleles gives *D*\* an advantage to detect locally introgressed regions by including more informative sites. When the true ancestral allele is known for all sites and single lineages are sampled from each population, the genome-wide power for *D*\*/*D*^+^ is less than *D*. However, if multiple lineages are sampled from each population, we have evidence to believe that *D*\* outperforms *D* regardless of polarization for genome-wide introgression detection in a variety of scenarios. This is of particular interest since outgroup samples are not obtainable, but also, an ‘optimal’ outgroup may not exist. A recent theoretical dissection of Patterson’s *D* demonstrated that a more distant outgroup can give rise to spurious signals of introgression Pang et al. (2025).

We evaluated *D*\*’s performance for detecting genome-wide introgression. In the case of large deviations in drift between the two sister populations, *D*\* exhibited a bias genome-wide. We showed that altering the standardized sample size can overcome this genome-wide bias. Treating the standardized sample size as a tuning parameter is an indirect way of incorporating variants of low frequency caused by intense drift in either population. This remedy needs to be further studied to confidently detect genome-wide introgression with *D*\*. For now, we do not recommend using *D*\* to detect genome-wide introgression since unequal drift between populations is expected in real populations. We suggest that a future extension of *D*\* is to correct for the case of strong drift and genome-wide introgression.

*D*\* provides a higher degree of certainty on outlier regions compared to *hmmix* and *Sprime*. When proposing candidate regions of introgression, it is desirable to form a consensus where several methods agree on proposed candidates Skov et al. (2018). This is particularly applicable to the pairing of *hmmix* and *Sprime* with *D*\* since *hmmix* and *Sprime* do not distinguish between different archaic populations and *hmmix* must be applied to each sample in the admixed population separately. *D*\* can distinguish between different archaic populations (Neanderthals or Denisovans) and groups all samples within each population together. To determine the archaic source of introgression with *hmmix* and *Sprime*, an additional analysis is required by matching candidate introgression regions with reference variation from different archaic populations.

We evaluated *D*\* on empirical samples by locating candidate regions of introgression between modern-day Papuans and Denisovans. We used *D*\* as an exploratory method on additional lineages from Papuans and found genes such as *RRM* 1, *HIC*2, *BANK*1, *CLEC*9*A*, and *LRBA*, which all have functions related to the immune system. This is consistent with previous analyses Jacobs et al. (2019); Vespasiani et al. (2022). In addition, we were able to recover regions belonging to *ADK*, a gene suspected for adaptation to high altitude González-Buenfil et al. (2024). We downsampled the number of Papuan individuals to a single diploid sample and showed that *D*\* when compared to *D*^+^ maintained a greater proportion of blocks found when the full set of Papuan samples were used.

There are some points of consideration when using *D*\*. The first is common to many methods relying on a block (window) size. This is largely dependent on recombination rate and the estimated age of the potentially introgressed segments Durvasula and Sankararaman (2019). Without estimated divergence times between populations, this can be difficult. We suggest running *D*\* multiple times with varying block sizes and looking for consistency in outlier regions. Finally, *D*\* and *D*^+^ provide an exploratory means of locating introgressed regions and lack measures of statistical significance in the absence of an underlying demographic model. Therefore, outlier regions identified with *D*\* should be labeled as candidates for further analysis.

## Conflicts of interest

The authors declare that they have no competing interests.

## Funding

This work was supported by the National Institute of General Medical Sciences of the National Institutes of Health award number R35GM146926 (Z.A.S.).

## Data availability

Precompiled executables, the manual, source code, and simulation and analysis scripts used in this study can be found at https://github.com/TQ-Smith/DSTAR. The Allen Ancient DNA Resource (AADR) can be accessed at https://dataverse.harvard.edu/dataset.xhtml?persistentId=doi:10.7910/DVN/FFIDCW. The Human Genome Diversity Project (HGDP) can be accessed at https://www.internationalgenome.org/data-portal/data-collection/HGDP.

## Author contributions statement

T.Q.S. and Z.A.S. conceived the experiment(s), T.Q.S. conducted the experiment(s), T.Q.S. analysed the results. T.Q.S. and Z.A.S. wrote and reviewed the manuscript.

## Acknowledgments

The authors would like to thank two anonymous reviewers. Computations for this research were performed using the Pennsylvania State University’s Institute for Computational Data Sciences’ Roar supercomputer.

## Notes

https://gnomad.broadinstitute.org/downloads

https://dataverse.harvard.edu/dataverse/reich_lab

### Competing Interest Statement

The authors have declared no competing interest.

### Summary of Updates

Corrected small in-line edits, namely captions.

## References

N. H. Barton and M. Slatkin. A quasi-equilibrium theory of the distribution of rare alleles in a subdivided population. Heredity, 56(3):409–415, 1986.

D. Y. Brandt, C. D. Huber, C. W. Chiang, and D. Ortega-Del Vecchyo. The promise of inferring the past using the ancestral recombination graph. Genome Biology and Evolution, 16(2):evae005, 2024.

S. R. Browning, B. L. Browning, Y. Zhou, S. Tucci, and J. M. Akey. Analysis of human sequence data reveals two pulses of archaic denisovan admixture. Cell, 173(1):53–61, 2018.

J. A. Cahill, P. D. Heintzman, K. Harris, M. D. Teasdale, J. Kapp, A. E. Soares, I. Stirling, D. Bradley, C. J. Edwards, K. Graim, et al. Genomic evidence of widespread admixture from polar bears into brown bears during the last ice age. Molecular Biology and Evolution, 35(5):1120–1129, 2018.

L. Chen, A. B. Wolf, W. Fu, L. Li, and J. M. Akey. Identifying and interpreting apparent neanderthal ancestry in african individuals. Cell, 180(4):677–687, 2020.

K. A. Cobb and M. L. Smith. The reasonable effectiveness of domain adaptation for inference of introgression. bioRxiv, pages 2025–01, 2025.

J. Dabney, M. Meyer, and S. Pääbo. Ancient dna damage. Cold Spring Harbor perspectives in biology, 5(7):a012567, 2013.

P. Danecek, J. K. Bonfield, J. Liddle, J. Marshall, V. Ohan, M. O. Pollard, A. Whitwham, T. Keane, S. A. McCarthy, R. M. Davies, et al. Twelve years of samtools and bcftools. Gigascience, 10(2):giab008, 2021.

M. De Manuel, M. Kuhlwilm, P. Frandsen, V. C. Sousa, T. Desai, J. Prado-Martinez, J. Hernandez-Rodriguez, I. Dupanloup, O. Lao, P. Hallast, et al. Chimpanzee genomic diversity reveals ancient admixture with bonobos. Science, 354(6311):477–481, 2016.

E. Y. Durand, N. Patterson, D. Reich, and M. Slatkin. Testing for ancient admixture between closely related populations. Molecular biology and evolution, 28(8):2239–2252, 2011.

A. Durvasula and S. Sankararaman. A statistical model for reference-free inference of archaic local ancestry. PLoS genetics, 15(5):e1008175, 2019.

A. Durvasula and S. Sankararaman. Recovering signals of ghost archaic introgression in african populations. Science Advances, 6(7):eaax5097, 2020.

N. B. Edelman and J. Mallet. Prevalence and adaptive impact of introgression. Annual review of genetics, 55(1):265–283, 2021.

N. B. Edelman, P. B. Frandsen, M. Miyagi, B. Clavijo, J. Davey, R. B. Dikow, G. García-Accinelli, S. M. Van Belleghem, N. Patterson, D. E. Neafsey, et al. Genomic architecture and introgression shape a butterfly radiation. Science, 366(6465): 594–599, 2019.

J. H. Gillespie. Population genetics: a concise guide. JHU press, 2004.

G. Gómez Hernández, M. Morell, and M. E. Alarcón-Riquelme. The role of bank1 in b cell signaling and disease. Cells, 10(5): 1184, 2021.

R. González-Buenfil, S. Vieyra-Sánchez, C. D. Quinto-Cortés, S. J. Oppenheimer, W. Pomat, M. Laman, M. C. Cervantes-Hernández, C. Barberena-Jonas, K. Auckland, A. Allen, et al. Genetic signatures of positive selection in human populations adapted to high altitude in papua new guinea. Genome biology and evolution, 16(8):evae161, 2024.

R. E. Green, J. Krause, A. W. Briggs, T. Maricic, U. Stenzel, M. Kircher, N. Patterson, H. Li, W. Zhai, M. H.-Y. Fritz, et al. A draft sequence of the neandertal genome. science, 328(5979): 710–722, 2010.

R. N. Gutenkunst, R. D. Hernandez, S. H. Williamson, and C. D. Bustamante. Inferring the joint demographic history of multiple populations from multidimensional snp frequency data. PLoS genetics, 5(10):e1000695, 2009.

M. W. Hahn and M. S. Hibbins. A three-sample test for introgression. Molecular biology and evolution, 36(12):2878–2882, 2019.

É. Harney, N. Patterson, D. Reich, and J. Wakeley. Assessing the performance of qpadm: a statistical tool for studying population admixture. Genetics, 217(4):iyaa045, 2021.

J. Hey. Isolation with migration models for more than two populations. Molecular biology and evolution, 27(4):905–920, 2010.

M. S. Hibbins and M. W. Hahn. Phylogenomic approaches to detecting and characterizing introgression. Genetics, 220(2): iyab173, 2022.

A. Hodgkinson and A. Eyre-Walker. Variation in the mutation rate across mammalian genomes. Nature reviews genetics, 12 (11):756–766, 2011.

P. Huang, S. A. Peslak, R. Ren, E. Khandros, K. Qin, C. A. Keller, B. Giardine, H. W. Bell, X. Lan, M. Sharma, et al. Hic2 controls developmental hemoglobin switching by repressing bcl11a transcription. Nature genetics, 54(9):1417–1426, 2022.

M. J. Hubisz, A. L. Williams, and A. Siepel. Mapping gene flow between ancient hominins through demography-aware inference of the ancestral recombination graph. PLoS genetics, 16(8): e1008895, 2020.

S. H. Hurlbert. The nonconcept of species diversity: a critique and alternative parameters. Ecology, 52(4):577–586, 1971.

G. S. Jacobs, G. Hudjashov, L. Saag, P. Kusuma, C. C. Darusallam, D. J. Lawson, M. Mondal, L. Pagani, F.-X. Ricaut, M. Stoneking, et al. Multiple deeply divergent denisovan ancestries in papuans. Cell, 177(4):1010–1021, 2019.

T. G. Jakobsson and Mattias. Population genomic analyses of dna from ancient remains. Handbook of Statistical Genomics: Two Volume Set, pages 295–40, 2019.

S. T. Kalinowski. Counting alleles with rarefaction: private alleles and hierarchical sampling designs. Conservation genetics, 5 (4):539–543, 2004.

J. Kamm, J. Terhorst, R. Durbin, and Y. S. Song. Efficiently inferring the demographic history of many populations with allele count data. Journal of the American Statistical Association, 115(531):1472–1487, 2020.

J. Kelleher, A. M. Etheridge, and G. McVean. Efficient coalescent simulation and genealogical analysis for large sample sizes. PLoS computational biology, 12(5):e1004842, 2016.

Z. Koenig, M. T. Yohannes, L. L. Nkambule, X. Zhao, J. K. Goodrich, H. A. Kim, M. W. Wilson, G. Tiao, S. P. Hao, N. Sahakian, et al. A harmonized public resource of deeply sequenced diverse human genomes. Genome Research, 34(5): 796–809, 2024.

D. J. Lawson, G. Hellenthal, S. Myers, and D. Falush. Inference of population structure using dense haplotype data. PLoS genetics, 8(1):e1002453, 2012.

L. Li and J. M. Akey. Statistical methods to detect archaic admixture and identify introgressed sequences. Handbook of Statistical Genomics: Two Volume Set, pages 275–20, 2019.

L. Li, T. J. Comi, R. F. Bierman, and J. M. Akey. Recurrent gene flow between neanderthals and modern humans over the past 200,000 years. Science, 385(6705):eadi1768, 2024.

L. Lopez Fang, D. Peede, D. Ortega-Del Vecchyo, E. J. McTavish, and E. Huerta-Sánchez. Leveraging shared ancestral variation to detect local introgression. PLoS Genetics, 20(1):e1010155, 2024.

G. Lopez-Herrera, G. Tampella, Q. Pan-Hammarström, P. Herholz, C. M. Trujillo-Vargas, K. Phadwal, A. K. Simon, M. Moutschen, A. Etzioni, A. Mory, et al. Deleterious mutations in lrba are associated with a syndrome of immune deficiency and autoimmunity. The American Journal of Human Genetics, 90(6):986–1001, 2012.

M. C. Macià, L. Skov, Z. E. Damgaard Bæk, and A. Hobolth. Enhancement of hidden markov model analyses for improved inference of archaic introgression in modern humans. bioRxiv, pages 2025–04, 2025.

C. Macià, L. Skov, Z. E. D. Bæk, and A. Hobolth. Enhancement of hidden markov model analyses for improved inference of archaic introgression in modern humans. Molecular Biology and Evolution, page msag134, 2026.

A.-S. Malaspinas, M. C. Westaway, C. Muller, V. C. Sousa, O. Lao Alves, A. Bergström, G. Athanasiadis, J. Y. Cheng, J. E. Crawford, et al. A genomic history of aboriginal australia. Nature, 538(7624):207–214, 2016.

M. Malinsky, M. Matschiner, and H. Svardal. Dsuite-fast d-statistics and related admixture evidence from vcf files. Molecular ecology resources, 21(2):584–595, 2021.

Mallick, A. Micco, M. Mah, H. Ringbauer, I. Lazaridis, I. Olalde, N. Patterson, and D. Reich. The allen ancient dna resource (aadr) a curated compendium of ancient human genomes. Scientific Data, 11(1):182, 2024.

S. H. Martin, J. W. Davey, and C. D. Jiggins. Evaluating the use of abba–baba statistics to locate introgressed loci. Molecular biology and evolution, 32(1):244–257, 2015.

M. Meyer, M. Kircher, M.-T. Gansauge, H. Li, F. Racimo, S. Mallick, J. G. Schraiber, F. Jay, K. Prüfer, C. De Filippo, et al. A high-coverage genome sequence from an archaic denisovan individual. science, 338(6104):222–226, 2012.

P. Moorjani, N. Patterson, J. N. Hirschhorn, A. Keinan, L. Hao,G. Atzmon, E. Burns, H. Ostrer, A. L. Price, and D. Reich. The history of african gene flow into southern europeans, levantines, and jews. PLoS genetics, 7(4):e1001373, 2011.

R. Nielsen, A. H. Vaughn, and Y. Deng. Inference and applications of ancestral recombination graphs. Nature Reviews Genetics, 26(1):47–58, 2025.

D. Pandey, M. Harris, N. R. Garud, and V. M. Narasimhan. Leveraging ancient dna to uncover signals of natural selection in europe lost due to admixture or drift. Nature Communications, 15(1):9772, 2024.

X.-X. Pang, J. Liu, and D.-Y. Zhang. Detecting introgression in shallow phylogenies: how minor molecular clock deviations lead to major inference errors. Molecular Biology and Evolution, 42(10):msaf216, 2025.

N. Patterson, P. Moorjani, Y. Luo, S. Mallick, N. Rohland, Y. Zhan, T. Genschoreck, T. Webster, and D. Reich. Ancient admixture in human history. Genetics, 192(3):1065–1093, 2012.

U. Peter-Okaka and D. Boison. Adenosine kinase: an epigenetic modulator and drug target. Journal of Inherited Metabolic Disease, 48(3):e70033, 2025.

V. Plagnol and J. D. Wall. Possible ancestral structure in human populations. PLoS genetics, 2(7):e105, 2006.

K. Prüfer, C. De Filippo, S. Grote, F. Mafessoni, P. Korlević, M. Hajdinjak, B. Vernot, L. Skov, P. Hsieh, S. Peyrégne, et al. A high-coverage neandertal genome from vindija cave in croatia. Science, 358(6363):655–658, 2017.

F. Racimo, D. Marnetto, and E. Huerta-Sánchez. Signatures of archaic adaptive introgression in present-day human populations. Molecular biology and evolution, 34(2):296–317, 2017.

D. D. Ray, L. Flagel, and D. R. Schrider. Introunet: identifying introgressed alleles via semantic segmentation. PLoS Genetics, 20(2):e1010657, 2024.

D. R. Schrider and A. D. Kern. Supervised machine learning for population genetics: a new paradigm. Trends in Genetics, 34 (4):301–312, 2018.

A. Sethuraman. The missing data problem in population genomics and statistical methods to address them, 2026.

L. Skov, R. Hui, V. Shchur, A. Hobolth, A. Scally, M. H. Schierup, and R. Durbin. Detecting archaic introgression using an unadmixed outgroup. PLoS genetics, 14(9):e1007641, 2018.

M. Slatkin. Rare alleles as indicators of gene flow. Evolution, 39 (1):53–65, 1985.

T. Q. Smith, A. Rahman, and Z. A. Szpiech. Eggs: Empirical genotype generalizer for samples. Bioinformatics Advances, page vbag125, 2026.

B. Sousa da Mota, S. Rubinacci, D. I. Cruz Dávalos, C. E. G. Amorim, M. Sikora, N. N. Johannsen, M. H. Szmyt, P. Włodarczak, A. Szczepanek, M. M. Przybyła, et al. Imputation of ancient human genomes. Nature Communications, 14(1):3660, 2023.

Z. A. Szpiech, M. Jakobsson, and N. A. Rosenberg. Adze: a rarefaction approach for counting alleles private to combinations of populations. Bioinformatics, 24(21):2498–2504, 2008.

N. Takahata and M. Slatkin. Private alleles in a partially isolated population ii. distribution of persistence time and probability of emigration. Theoretical population biology, 30(2):180–193, 1986.

B. Vernot and J. M. Akey. Resurrecting surviving neandertal lineages from modern human genomes. Science, 343(6174): 1017–1021, 2014.

D. M. Vespasiani, G. S. Jacobs, L. E. Cook, N. Brucato, M. Leavesley, C. Kinipi, F.-X. Ricaut, M. P. Cox, and I. G. Romero. Denisovan introgression has shaped the immune system of present-day papuans. PLoS genetics, 18(12): e1010470, 2022.

B. F. Voight, S. Kudaravalli, X. Wen, and J. K. Pritchard. A map of recent positive selection in the human genome. PLoS biology, 4(3):e72, 2006.

M. C. Whitlock and D. E. McCauley. Indirect measures of gene flow and migration: Fs t ≠ 1/(4nm+ 1). Heredity, 82(2):117–125, 1999.

K. Yuan, X. Ni, C. Liu, Y. Pan, L. Deng, R. Zhang, Y. Gao, X. Ge, J. Liu, X. Ma, et al. Refining models of archaic admixture in eurasia with archaicseeker 2.0. Nature communications, 12(1): 6232, 2021.

J.-G. Zhang, P. E. Czabotar, A. N. Policheni, I. Caminschi, S. San Wan, S. Kitsoulis, K. M. Tullett, A. Y. Robin, R. Brammananth, M. F. van Delft, et al. The dendritic cell receptor clec9a binds damaged cells via exposed actin filaments. Immunity, 36(4):646–657, 2012.

W. Zhang, K. K. Dasmahapatra, J. Mallet, G. R. Moreira, and M. R. Kronforst. Genome-wide introgression among distantly related heliconius butterfly species. Genome biology, 17(1):25, 2016.

Z. Zhou, Q. Song, Y. Yang, L. Wang, and Z. Wu. Comprehensive landscape of rrm2 with immune infiltration in pan-cancer. Cancers, 14(12):2938, 2022.

