## Supplemental for "A Rarefaction Approach to Identify Local Introgression in a Three Population Tree"

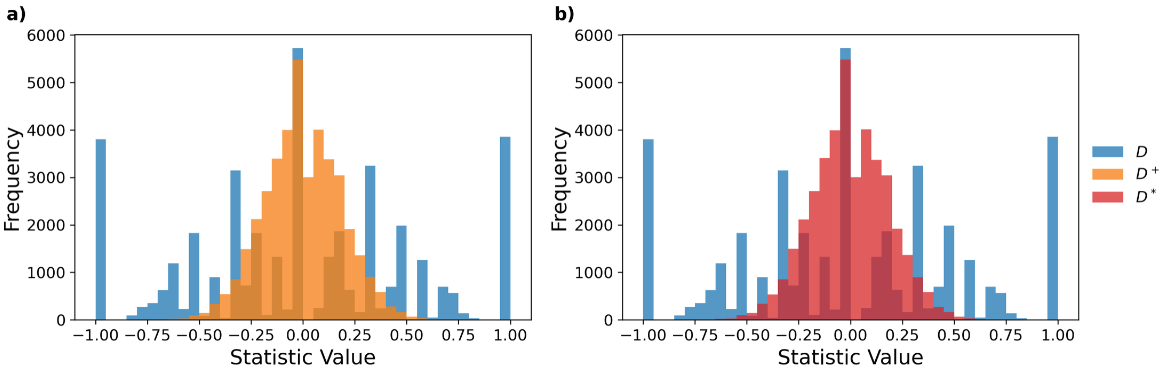


Figure S1: **Null Distribution for Single Lineages** **a)** The null-distribution of $D^+$ compared to $D$, and **b)** the null-distribution of $D^*$ compared to $D$ for single lineages.

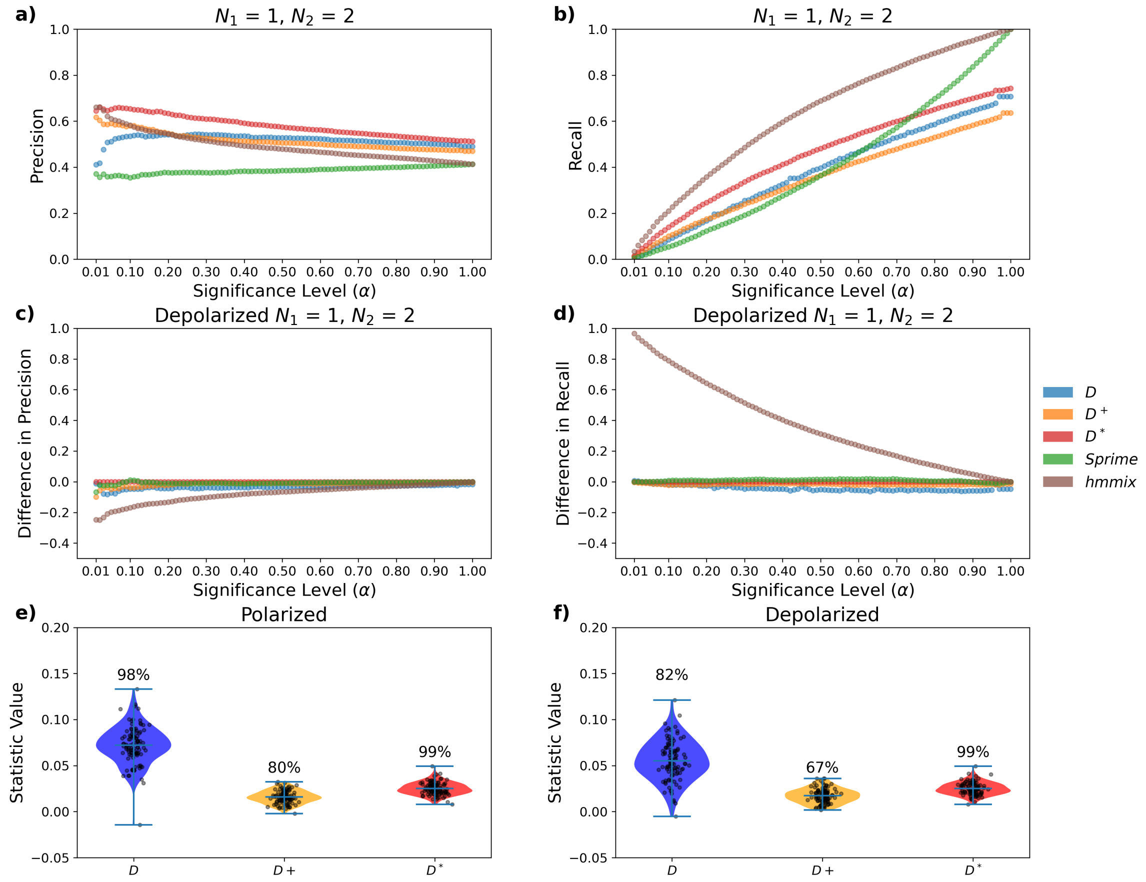


Figure S2: **Results for diploid sampling $P_1 = 1$ and $P_2 = 2$ a)** Precision and **b)** recall are shown for significance levels ($\alpha$) between 0.01 and 1. **c)** Difference (polarized minus depolarized) in precision and **d)** in recall. **e)** Genome-wide power for introgression detection on polarized genotypes, distribution gives local window values, percentage above gives power. **f)** Genome-wide power for introgression detection on unpolarized genotypes, distribution genome-wide values, percentage above gives power.

| | $N_2 = 1$ | $N_2 = 2$ | $N_2 = 6$ | $N_2 = 12$ |
|---|---|---|---|---|
| $N_1 = 1$ | $D$: −0.004/0.447<br>$D^+$: −0.000/0.137<br>$D^*$: −0.000/0.209 | $D$: 0.002/0.402<br>$D^+$: −0.002/0.124<br>$D^*$: −0.003/0.104 | $D$: 0.003/0.370<br>$D^+$: −0.004/0.115<br>$D^*$: −0.001/0.058 | $D$: 0.003/0.362<br>$D^+$: −0.004/0.113<br>$D^*$: −0.000/0.046 |
| $N_1 = 2$ | $D$: 0.001/0.401<br>$D^+$: 0.003/0.124<br>$D^*$: 0.003/0.104 | $D$: 0.001/0.355<br>$D^+$: 0.000/0.110<br>$D^*$: 0.000/0.063 | $D$: −0.003/0.321<br>$D^+$: −0.002/0.099<br>$D^*$: −0.001/0.038 | $D$: 0.002/0.313<br>$D^+$: −0.001/0.095<br>$D^*$: −0.000/0.030 |
| $N_1 = 6$ | $D$: −0.003/0.367<br>$D^+$: 0.002/0.114<br>$D^*$: 0.001/0.057 | $D$: −0.002/0.322<br>$D^+$: 0.001/0.099<br>$D^*$: 0.000/0.038 | $D$: −0.000/0.285<br>$D^+$: 0.000/0.086<br>$D^*$: −0.000/0.026 | $D$: 0.000/0.275<br>$D^+$: −0.001/0.083<br>$D^*$: 0.000/0.021 |
| $N_1 = 12$ | $D$: −0.004/0.364<br>$D^+$: 0.003/0.113<br>$D^*$: 0.001/0.046 | $D$: −0.002/0.312<br>$D^+$: 0.001/0.096<br>$D^*$: 0.000/0.030 | $D$: 0.002/0.276<br>$D^+$: 0.000/0.083<br>$D^*$: 0.000/0.021 | $D$: 0.000/0.267<br>$D^+$: −0.000/0.080<br>$D^*$: 0.000/0.018 |

Table S1: **Null-Distribution for Different Sample Sizes** The null distribution for $D$, $D^+$, and $D^*$ with different samples sizes. We do not show the mean and standard deviation for *Sprime* and *hmmix* since they are not expected to be centered around zero.

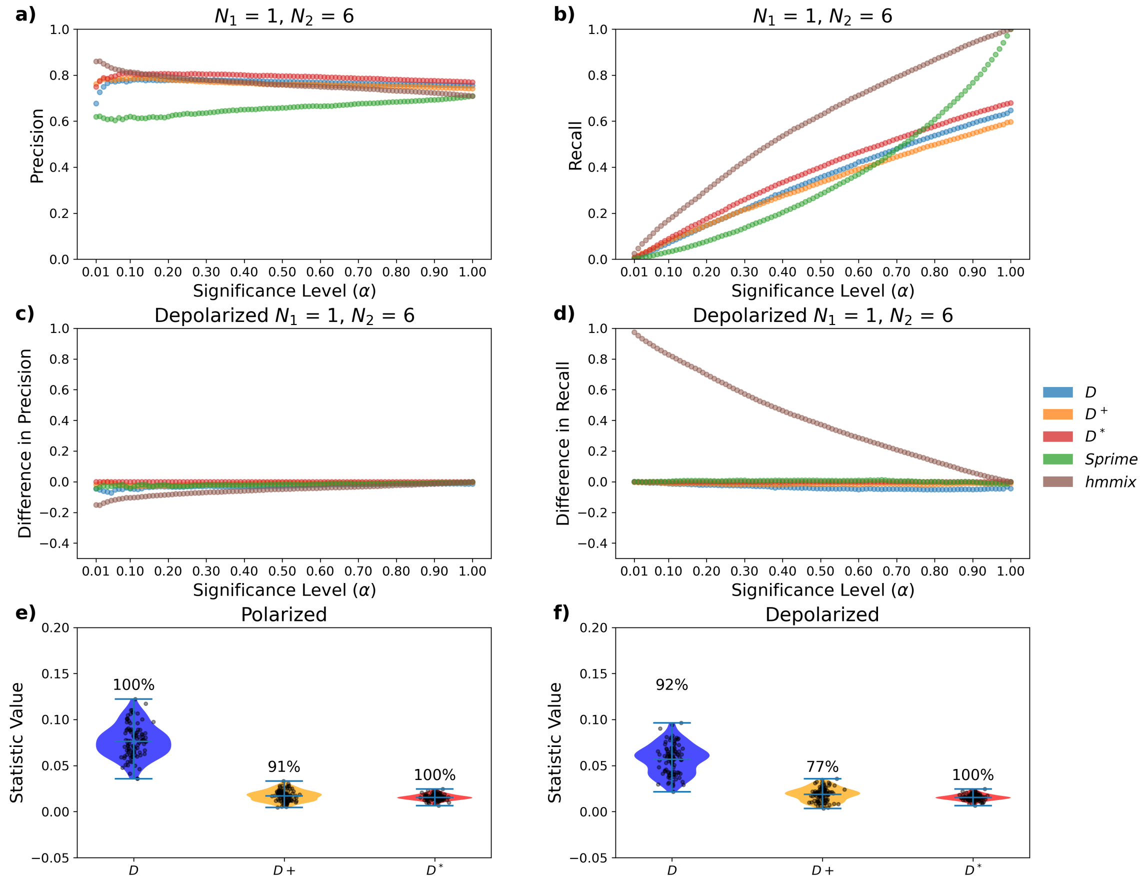


Figure S3: **Results for diploid sampling** $P_1 = 1$ **and** $P_2 = 6$ **a)** Precision and **b)** recall are shown for significance levels ($\alpha$) between 0.01 and 1. **c)** Difference (polarized minus depolarized) in precision and **d)** in recall. **e)** Genome-wide power for introgression detection on polarized genotypes, distribution gives local window values, percentage above gives power. **f)** Genome-wide power for introgression detection on unpolarized genotypes, distribution genome-wide values, percentage above gives power.

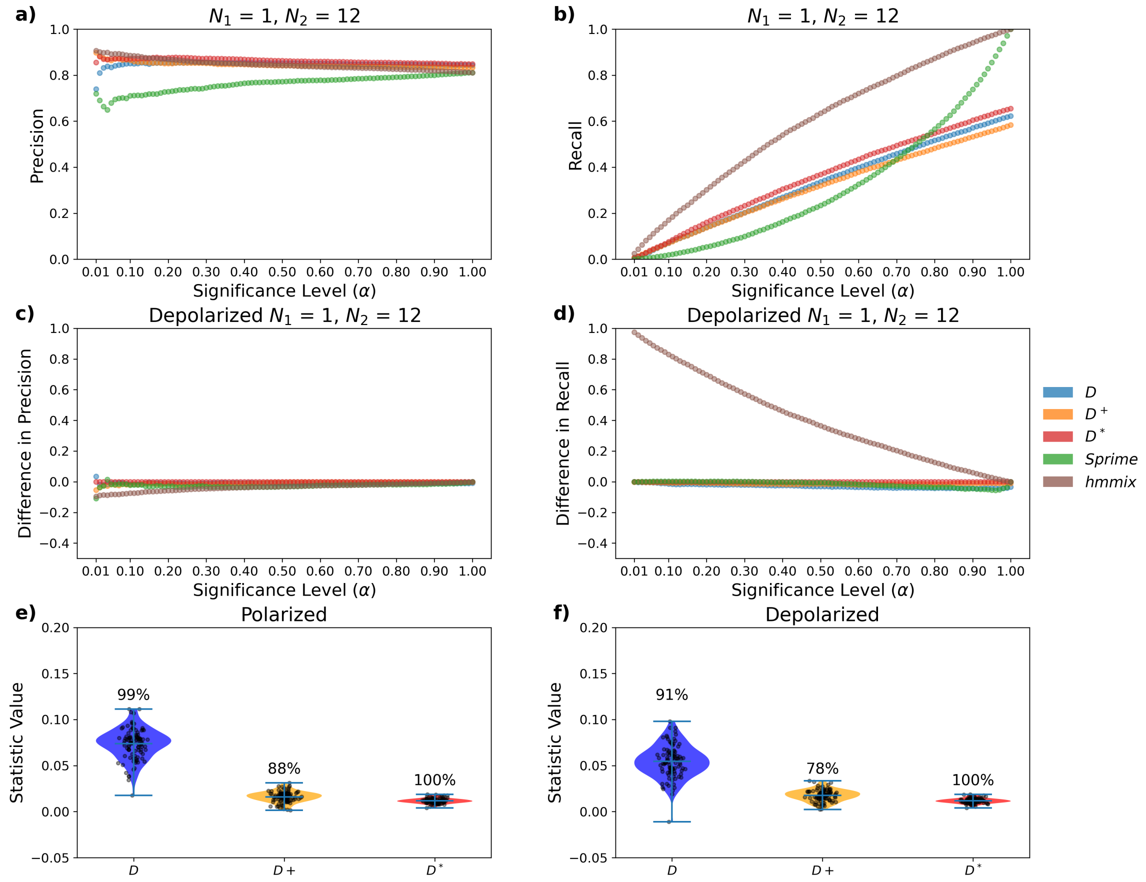


Figure S4: **Results for diploid sampling $P_1 = 1$ and $P_2 = 12$ a)** Precision and **b)** recall are shown for significance levels ($\alpha$) between 0.01 and 1. **c)** Difference (polarized minus depolarized) in precision and **d)** in recall. **e)** Genome-wide power for introgression detection on polarized genotypes, distribution gives local window values, percentage above gives power. **f)** Genome-wide power for introgression detection on unpolarized genotypes, distribution genome-wide values, percentage above gives power.

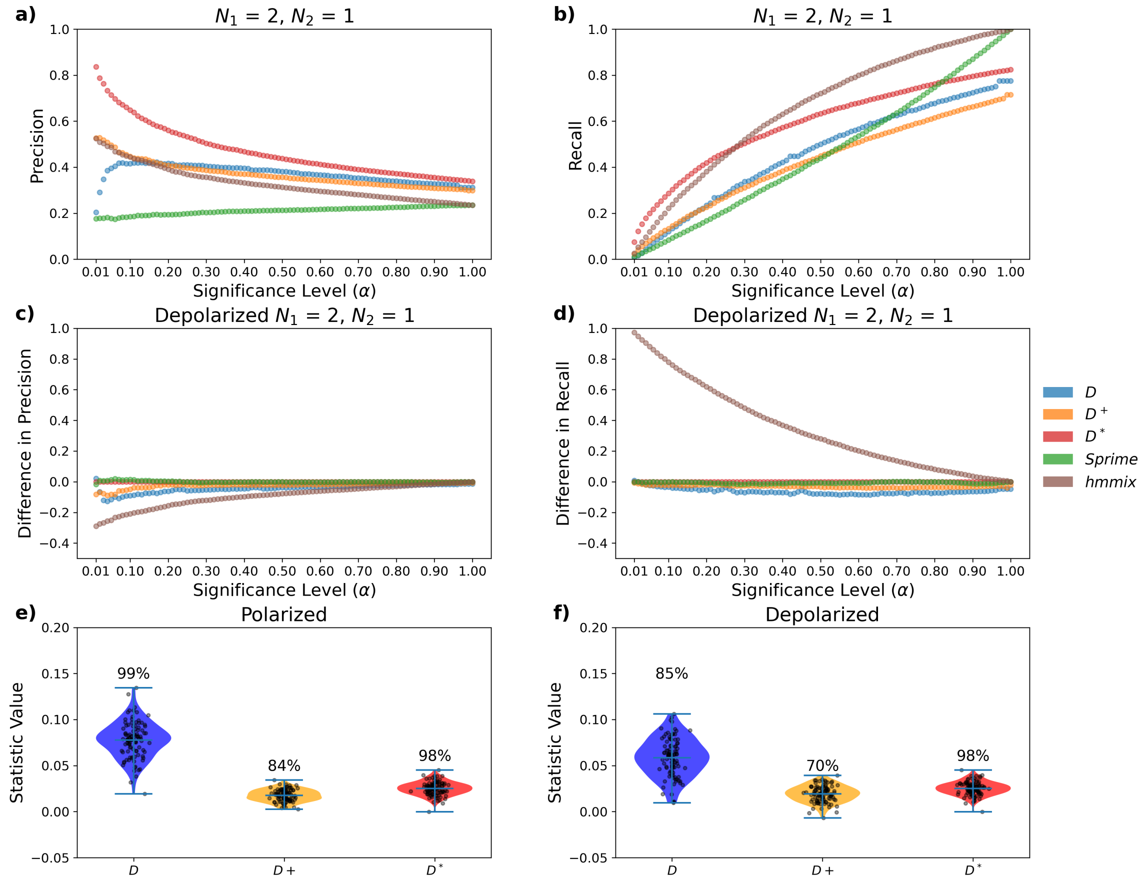


Figure S5: **Results for diploid sampling** $P_1 = 2$ **and** $P_2 = 1$ **a)** Precision and **b)** recall are shown for significance levels ($\alpha$) between 0.01 and 1. **c)** Difference (polarized minus depolarized) in precision and **d)** in recall. **e)** Genome-wide power for introgression detection on polarized genotypes, distribution gives local window values, percentage above gives power. **f)** Genome-wide power for introgression detection on unpolarized genotypes, distribution genome-wide values, percentage above gives power.

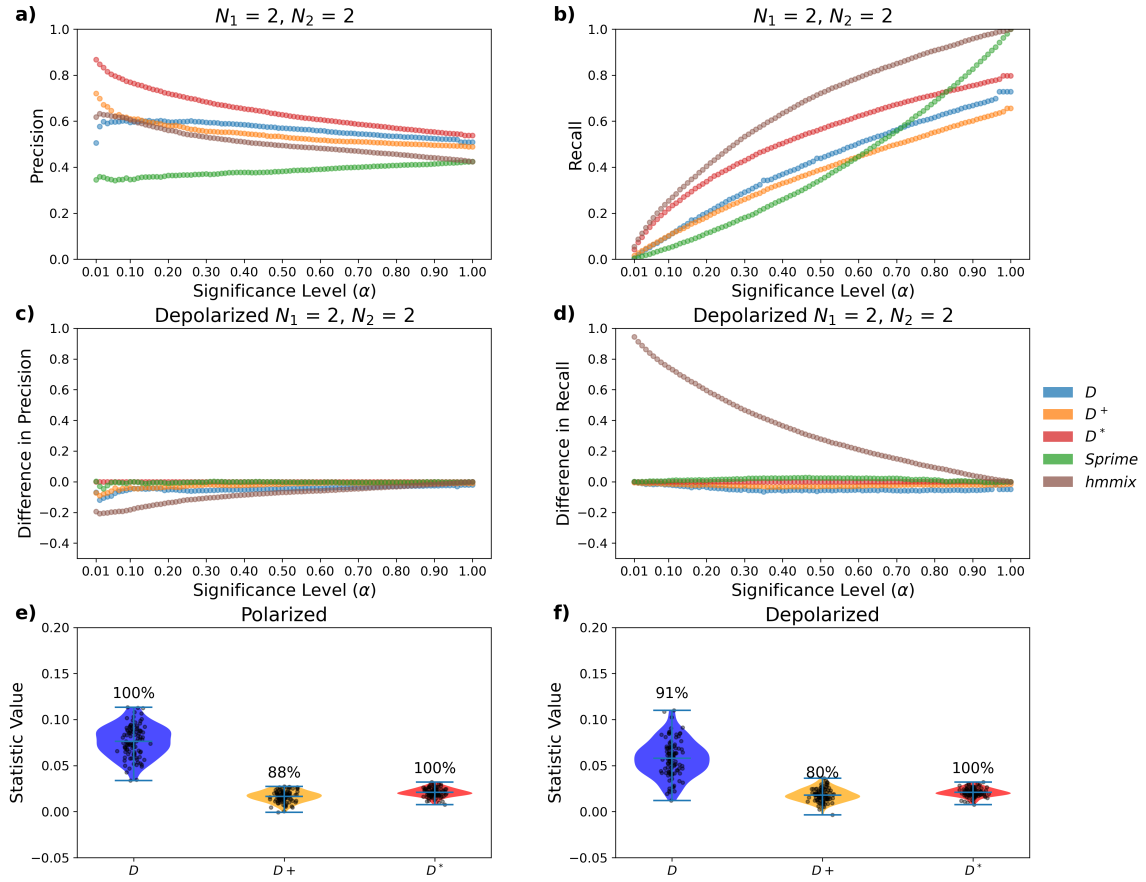


Figure S6: **Results for diploid sampling $P_1 = 2$ and $P_2 = 2$ a)** Precision and **b)** recall are shown for significance levels ($\alpha$) between 0.01 and 1. **c)** Difference (polarized minus depolarized) in precision and **d)** in recall. **e)** Genome-wide power for introgression detection on polarized genotypes, distribution gives local window values, percentage above gives power. **f)** Genome-wide power for introgression detection on unpolarized genotypes, distribution genome-wide values, percentage above gives power.

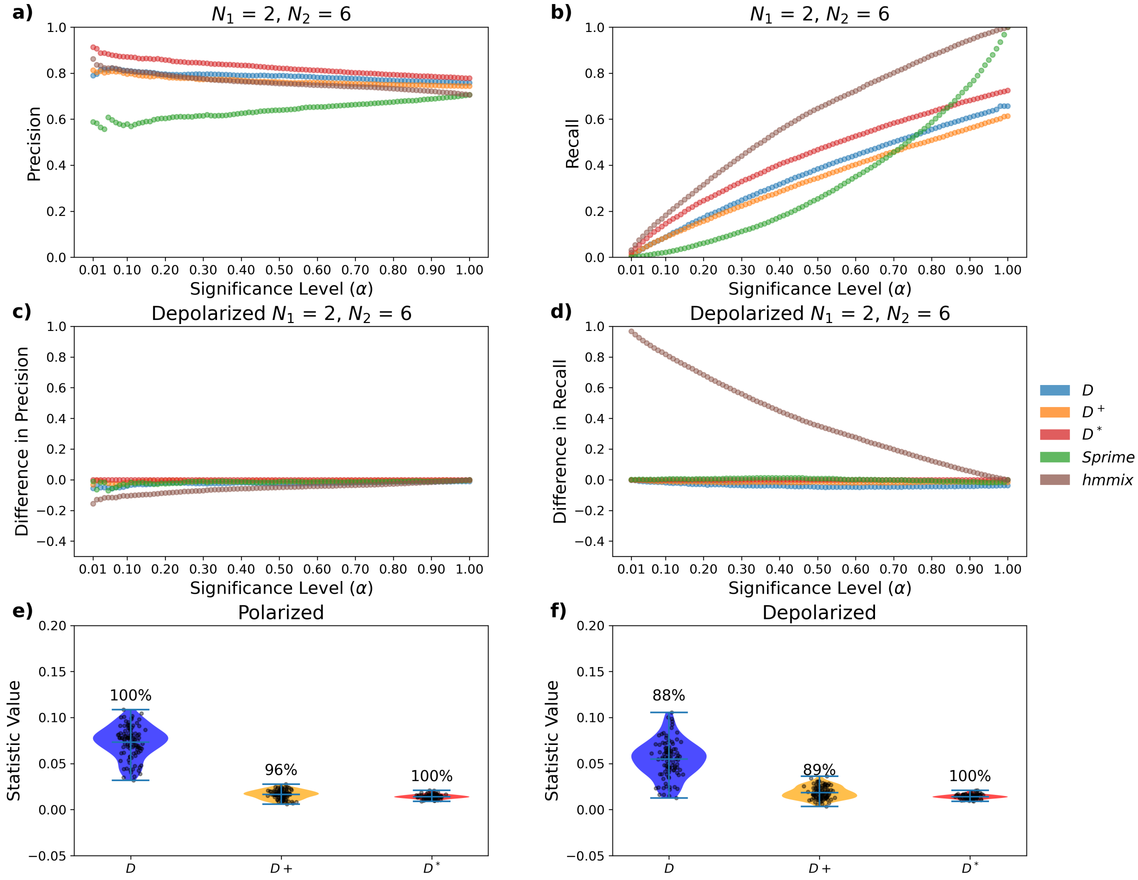


Figure S7: **Results for diploid sampling $P_1 = 2$ and $P_2 = 6$ a)** Precision and **b)** recall are shown for significance levels ($\alpha$) between 0.01 and 1. **c)** Difference (polarized minus depolarized) in precision and **d)** in recall. **e)** Genome-wide power for introgression detection on polarized genotypes, distribution gives local window values, percentage above gives power. **f)** Genome-wide power for introgression detection on unpolarized genotypes, distribution genome-wide values, percentage above gives power.

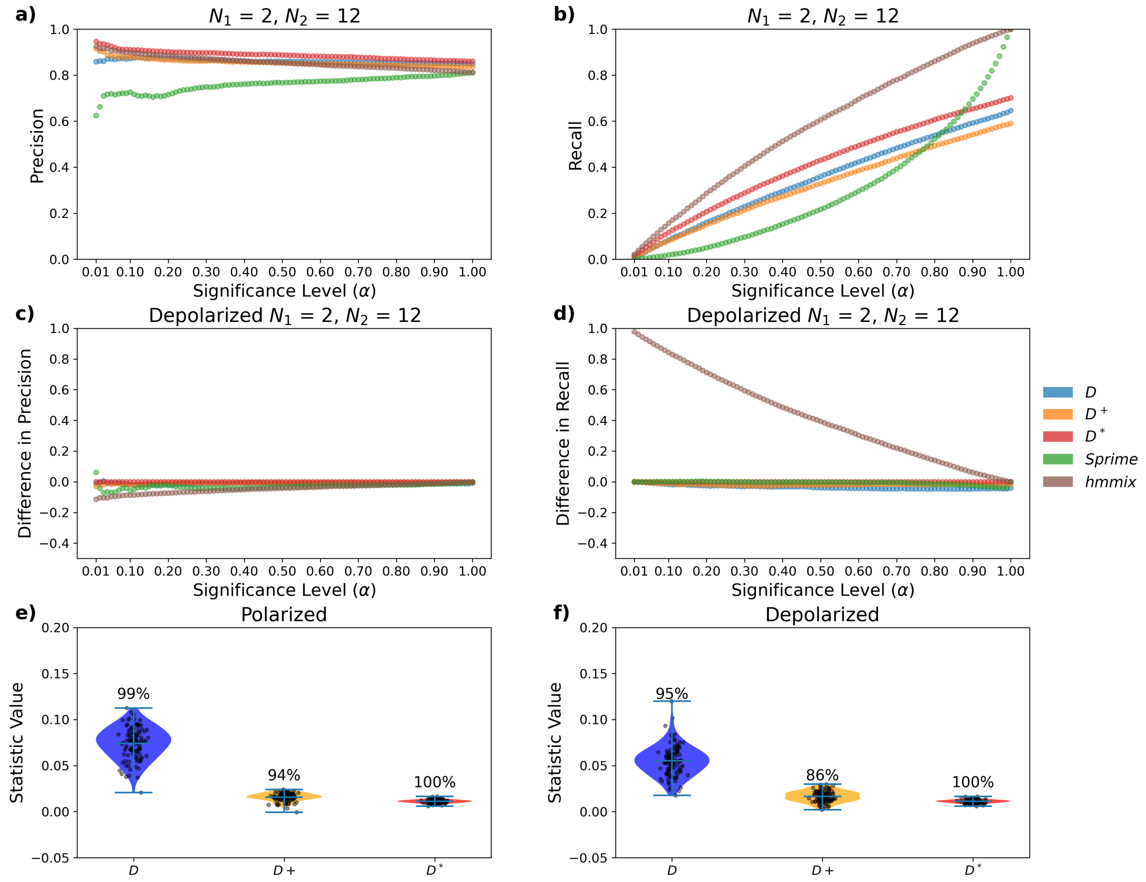


Figure S8: **Results for diploid sampling $P_1 = 2$ and $P_2 = 12$ a)** Precision and **b)** recall are shown for significance levels ($\alpha$) between 0.01 and 1. **c)** Difference (polarized minus depolarized) in precision and **d)** in recall. **e)** Genome-wide power for introgression detection on polarized genotypes, distribution gives local window values, percentage above gives power. **f)** Genome-wide power for introgression detection on unpolarized genotypes, distribution genome-wide values, percentage above gives power.

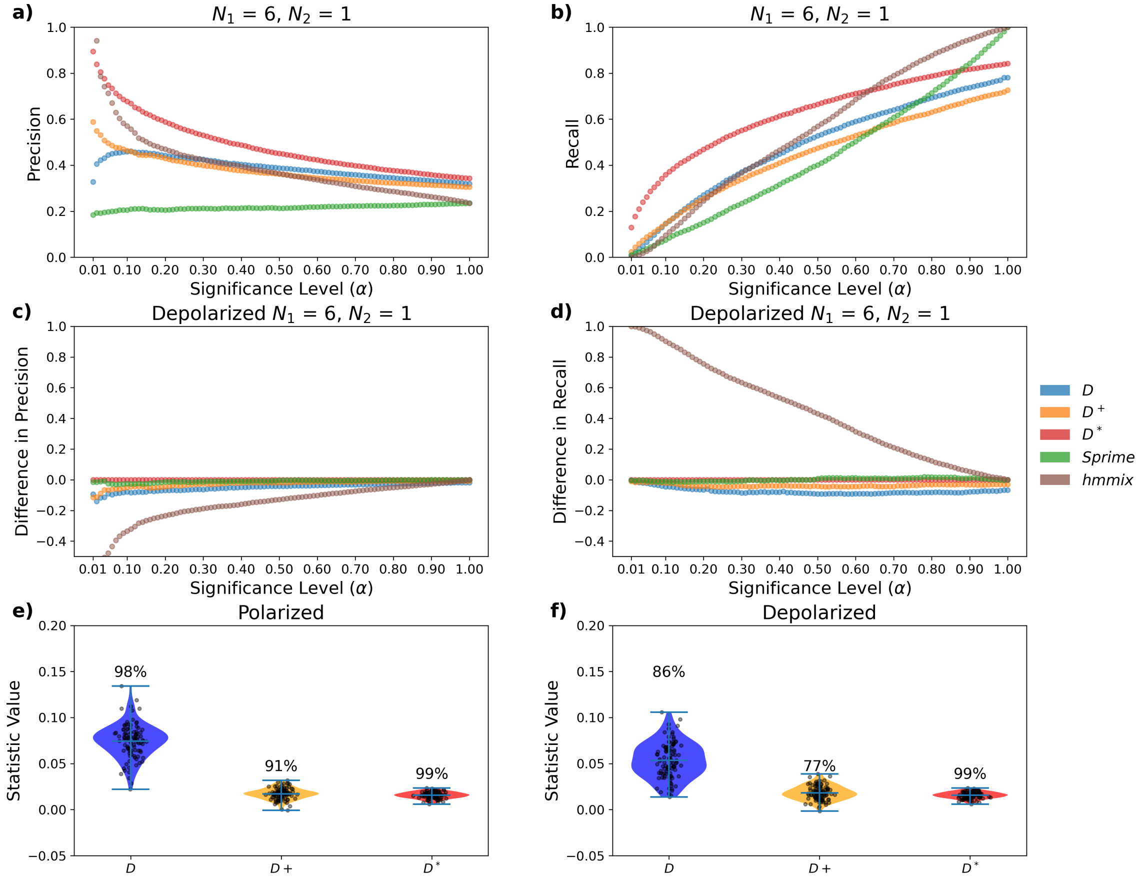


Figure S9: **Results for diploid sampling $P_1 = 6$ and $P_2 = 1$ a)** Precision and **b)** recall are shown for significance levels ($\alpha$) between 0.01 and 1. **c)** Difference (polarized minus depolarized) in precision and **d)** in recall. **e)** Genome-wide power for introgression detection on polarized genotypes, distribution gives local window values, percentage above gives power. **f)** Genome-wide power for introgression detection on unpolarized genotypes, distribution genome-wide values, percentage above gives power.

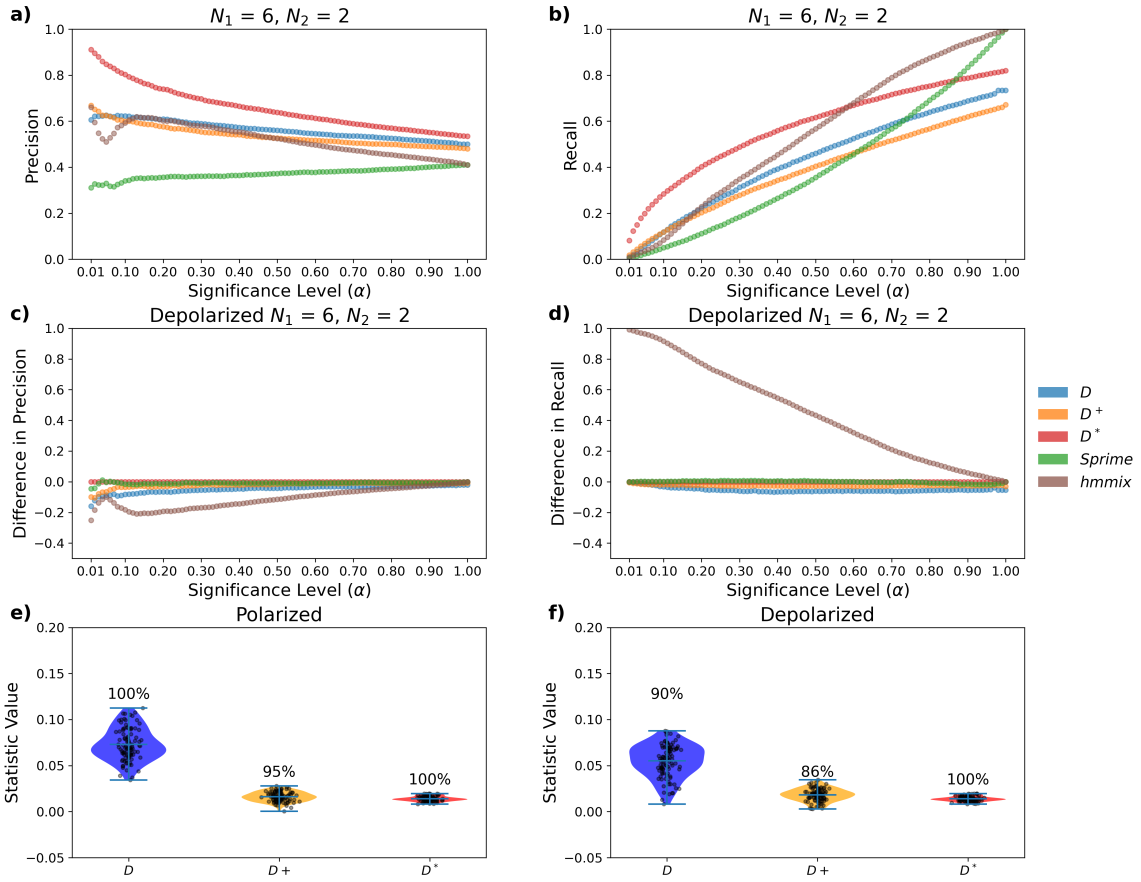


Figure S10: **Results for diploid sampling** $P_1 = 6$ **and** $P_2 = 2$ **a)** Precision and **b)** recall are shown for significance levels ($\alpha$) between 0.01 and 1. **c)** Difference (polarized minus depolarized) in precision and **d)** in recall. **e)** Genome-wide power for introgression detection on polarized genotypes, distribution gives local window values, percentage above gives power. **f)** Genome-wide power for introgression detection on unpolarized genotypes, distribution genome-wide values, percentage above gives power.

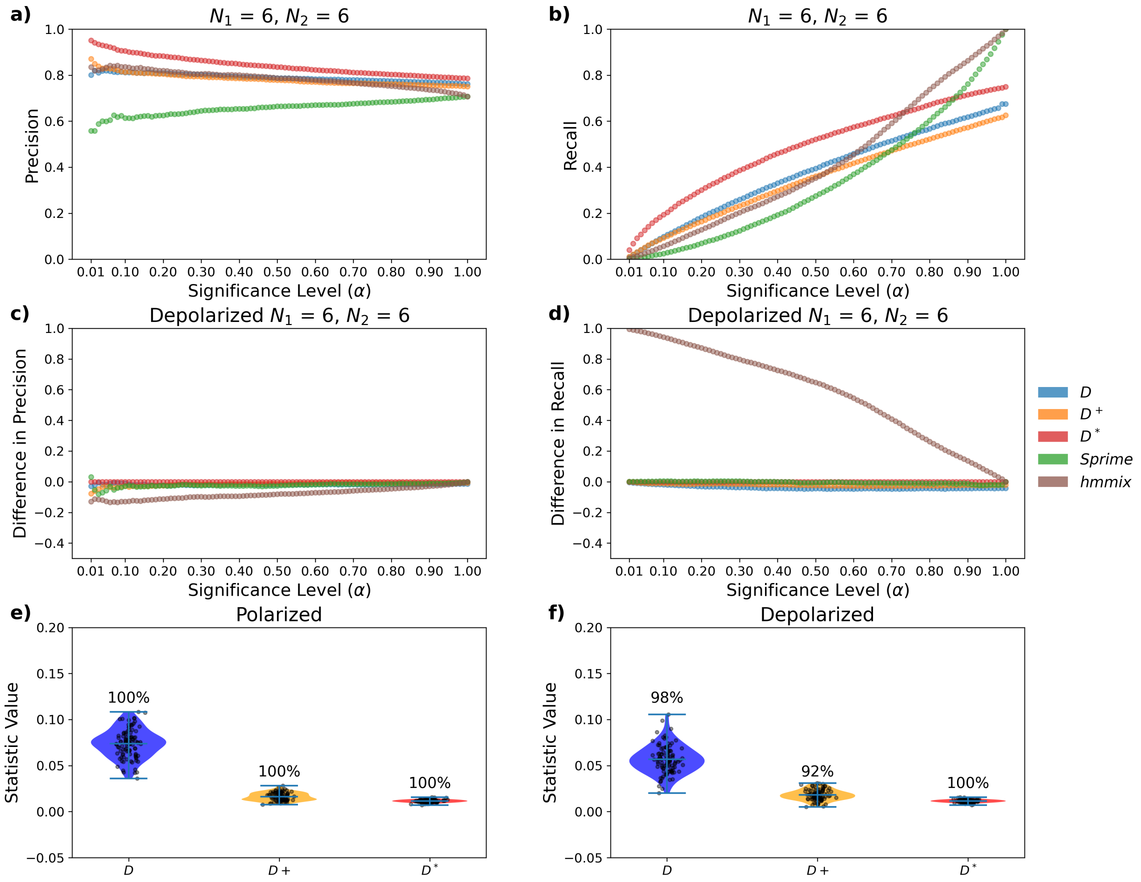


Figure S11: **Results for diploid sampling** $P_1 = 6$ **and** $P_2 = 6$ **a)** Precision and **b)** recall are shown for significance levels ($\alpha$) between 0.01 and 1. **c)** Difference (polarized minus depolarized) in precision and **d)** in recall. **e)** Genome-wide power for introgression detection on polarized genotypes, distribution gives local window values, percentage above gives power. **f)** Genome-wide power for introgression detection on unpolarized genotypes, distribution genome-wide values, percentage above gives power.

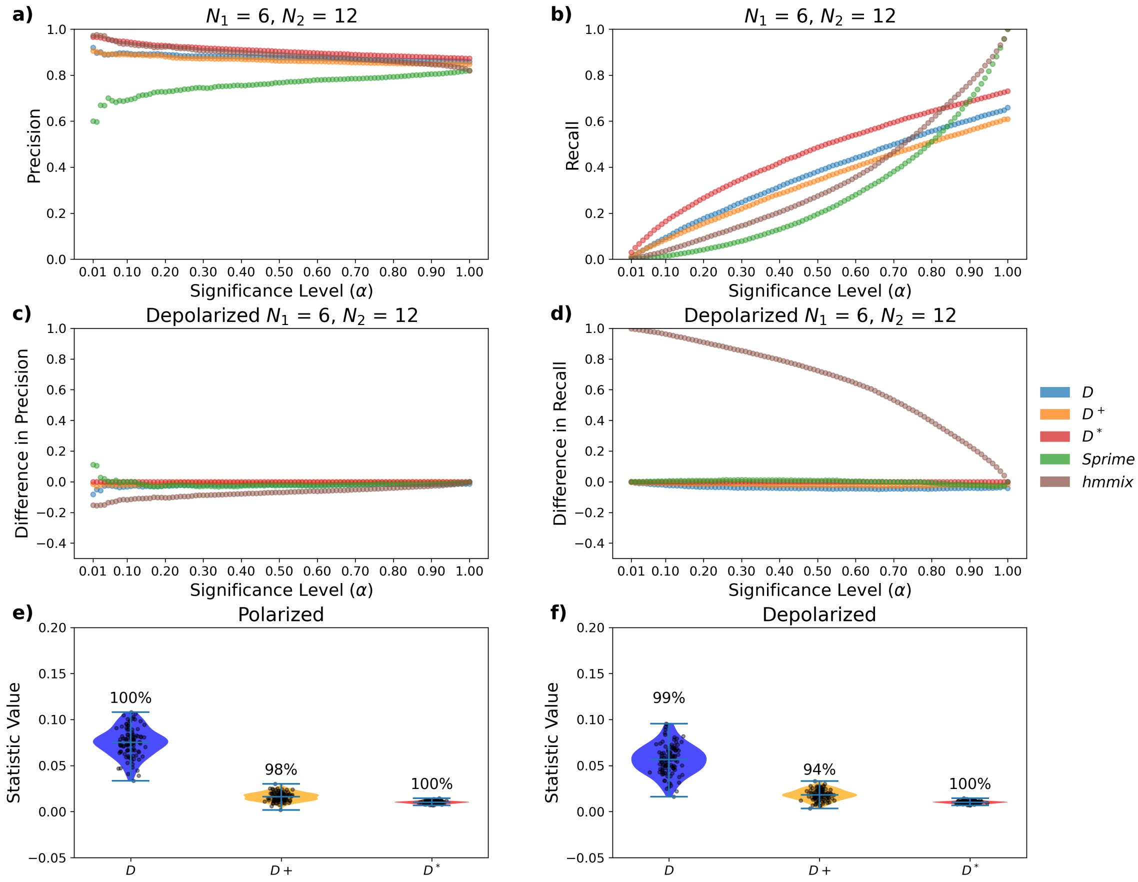


Figure S12: **Results for diploid sampling** $P_1 = 6$ **and** $P_2 = 12$ **a)** Precision and **b)** recall are shown for significance levels ($\alpha$) between 0.01 and 1. **c)** Difference (polarized minus depolarized) in precision and **d)** in recall. **e)** Genome-wide power for introgression detection on polarized genotypes, distribution gives local window values, percentage above gives power. **f)** Genome-wide power for introgression detection on unpolarized genotypes, distribution genome-wide values, percentage above gives power.

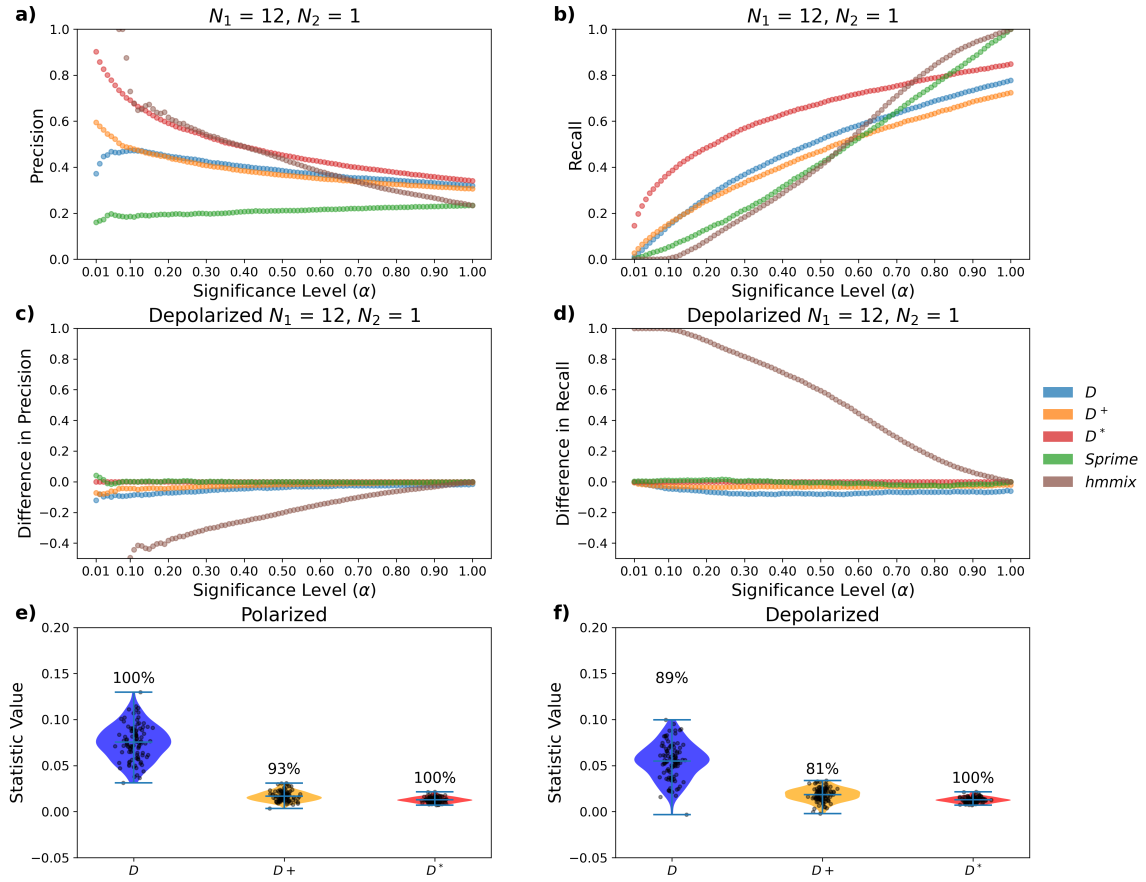


Figure S13: **Results for diploid sampling $P_1 = 12$ and $P_2 = 1$ a)** Precision and **b)** recall are shown for significance levels ($\alpha$) between 0.01 and 1. **c)** Difference (polarized minus depolarized) in precision and **d)** in recall. **e)** Genome-wide power for introgression detection on polarized genotypes, distribution gives local window values, percentage above gives power. **f)** Genome-wide power for introgression detection on unpolarized genotypes, distribution genome-wide values, percentage above gives power.

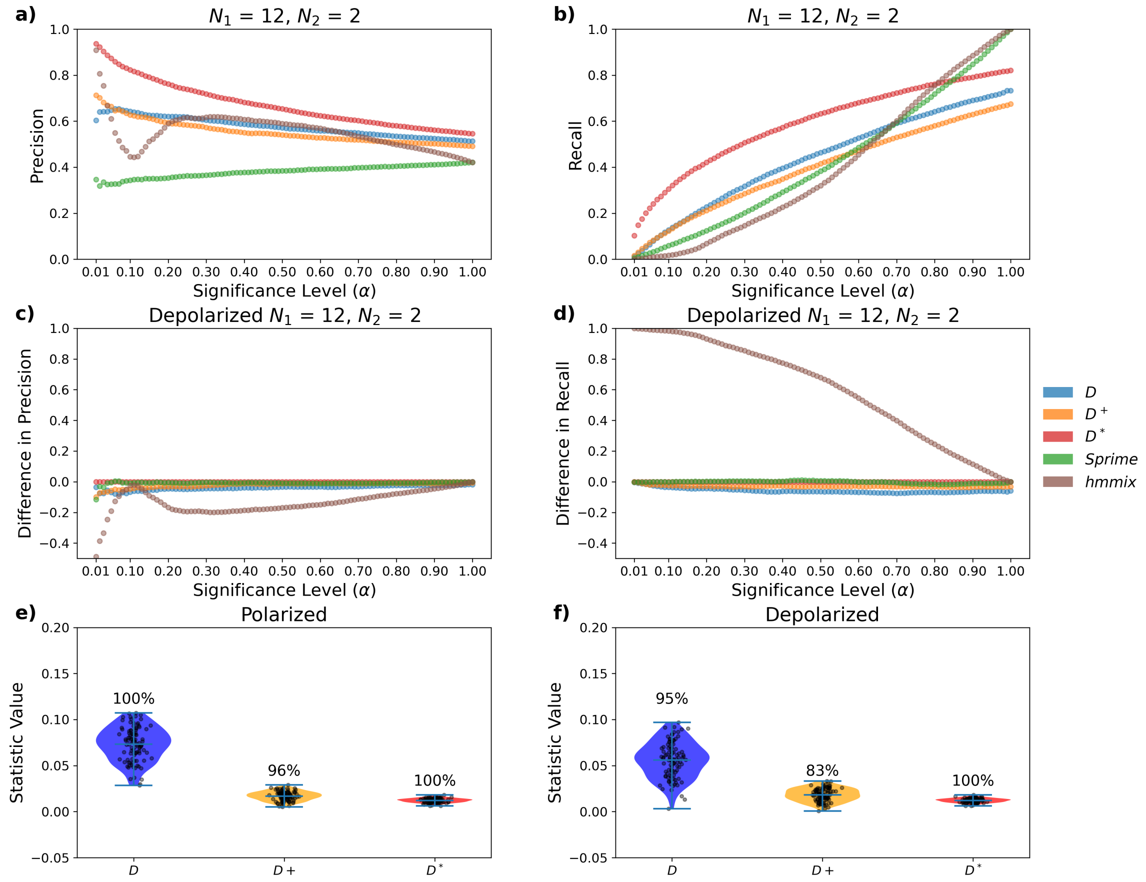


Figure S14: **Results for diploid sampling $P_1 = 12$ and $P_2 = 2$ a)** Precision and **b)** recall are shown for significance levels ($\alpha$) between 0.01 and 1. **c)** Difference (polarized minus depolarized) in precision and **d)** in recall. **e)** Genome-wide power for introgression detection on polarized genotypes, distribution gives local window values, percentage above gives power. **f)** Genome-wide power for introgression detection on unpolarized genotypes, distribution genome-wide values, percentage above gives power.

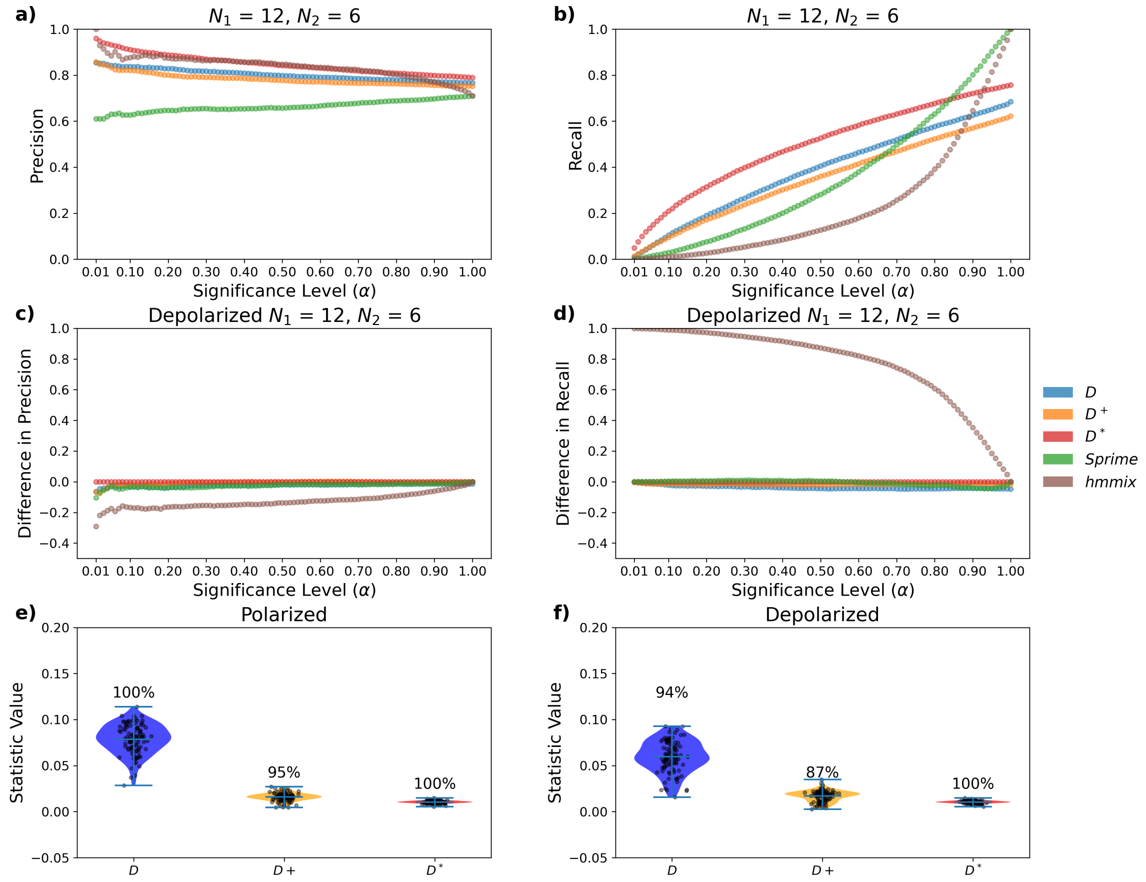


Figure S15: **Results for diploid sampling** $P_1 = 12$ **and** $P_2 = 6$ **a)** Precision and **b)** recall are shown for significance levels ($\alpha$) between 0.01 and 1. **c)** Difference (polarized minus depolarized) in precision and **d)** in recall. **e)** Genome-wide power for introgression detection on polarized genotypes, distribution gives local window values, percentage above gives power. **f)** Genome-wide power for introgression detection on unpolarized genotypes, distribution genome-wide values, percentage above gives power.

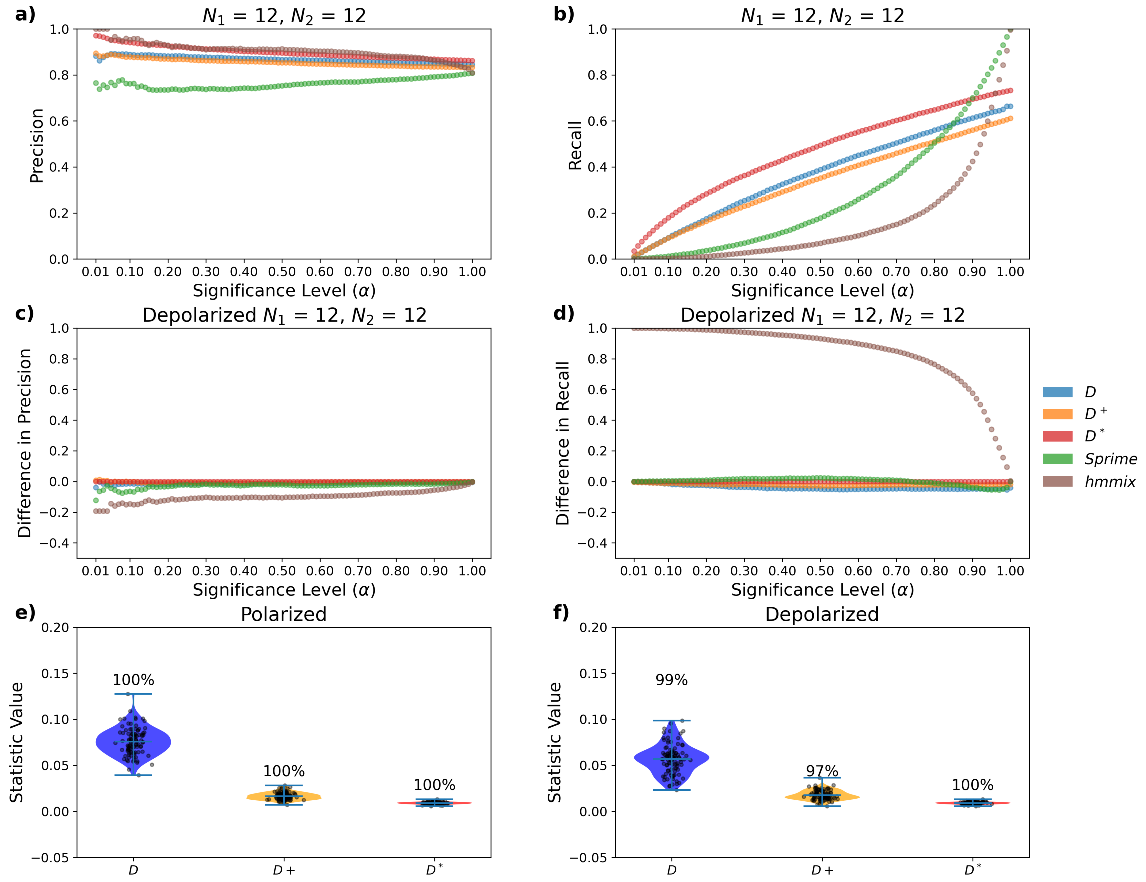


Figure S16: **Results for diploid sampling** $P_1 = 12$ **and** $P_2 = 12$ **a)** Precision and **b)** recall are shown for significance levels ($\alpha$) between 0.01 and 1. **c)** Difference (polarized minus depolarized) in precision and **d)** in recall. **e)** Genome-wide power for introgression detection on polarized genotypes, distribution gives local window values, percentage above gives power. **f)** Genome-wide power for introgression detection on unpolarized genotypes, distribution genome-wide values, percentage above gives power.

| | $N_2 = 1$ | $N_2 = 2$ | $N_2 = 6$ | $N_2 = 12$ |
|---|---|---|---|---|
| $N_1 = 1$ | $D$: $-18\%$<br>$D^+$: $-6\%$<br>$D^*$: $0\%$ | $D$: $-16\%$<br>$D^+$: $-13\%$<br>$D^*$: $0\%$ | $D$: $-8\%$<br>$D^+$: $-14\%$<br>$D^*$: $0\%$ | $D$: $-8\%$<br>$D^+$: $-10\%$<br>$D^*$: $0\%$ |
| $N_1 = 2$ | $D$: $-14\%$<br>$D^+$: $-14\%$<br>$D^*$: $0\%$ | $D$: $-9\%$<br>$D^+$: $-8\%$<br>$D^*$: $0\%$ | $D$: $-12\%$<br>$D^+$: $-7\%$<br>$D^*$: $0\%$ | $D$: $-4\%$<br>$D^+$: $-8\%$<br>$D^*$: $0\%$ |
| $N_1 = 6$ | $D$: $-12\%$<br>$D^+$: $-14\%$<br>$D^*$: $0\%$ | $D$: $-10\%$<br>$D^+$: $-9\%$<br>$D^*$: $0\%$ | $D$: $-2\%$<br>$D^+$: $-8\%$<br>$D^*$: $0\%$ | $D$: $-1\%$<br>$D^+$: $-4\%$<br>$D^*$: $0\%$ |
| $N_1 = 12$ | $D$: $-11\%$<br>$D^+$: $-12\%$<br>$D^*$: $0\%$ | $D$: $-5\%$<br>$D^+$: $-13\%$<br>$D^*$: $0\%$ | $D$: $-6\%$<br>$D^+$: $-8\%$<br>$D^*$: $0\%$ | $D$: $-1\%$<br>$D^+$: $-3\%$<br>$D^*$: $0\%$ |

Table S2: **Decrease in Genome-Wide Power Due to Depolarization** Change in genome-wide power between polarized and depolarized sites for each combination of $P_1 = \{1, 2, 6, 12\}$ and $P_2 = \{1, 2, 6, 12\}$.

| | $\lambda = 1.2$ | $\lambda = 1.5$ | $\lambda = 2$ | $\lambda = 5$ |
|---|---|---|---|---|
| P1 | $D$: $-0.002/0.449$<br>$D^+$: $0.019/0.137$<br>$D^*$: $0.020/0.205$ | $D$: $-0.000/0.452$<br>$D^+$: $0.045/0.136$<br>$D^*$: $0.047/0.201$ | $D$: $0.002/0.456$<br>$D^+$: $0.087/0.133$<br>$D^*$: $0.092/0.192$ | $D$: $0.001/0.484$<br>$D^+$: $0.277/0.121$<br>$D^*$: $0.275/0.149$ |
| P2 | $D$: $-0.002/0.448$<br>$D^+$: $-0.019/0.136$<br>$D^*$: $-0.017/0.207$ | $D$: $0.002/0.451$<br>$D^+$: $-0.046/0.136$<br>$D^*$: $-0.048/0.202$ | $D$: $0.000/0.459$<br>$D^+$: $-0.087/0.134$<br>$D^*$: $-0.091/0.193$ | $D$: $0.003/0.489$<br>$D^+$: $-0.276/0.121$<br>$D^*$: $-0.275/0.149$ |

Table S3: **Mean and Standard Deviation of Null-Distribution for Molecular Clock Violations** The mean and standard deviation of the null distribution for $D$, $D^+$, and $D^*$ with molecular clock violations. The mutation rate for each population is changed by a factor of $\lambda = \{1.2, 1.5, 2, 5\}$ with respect to the other population. We do not show the mean and standard deviation for *Sprime* and *hmmix* since they are not expected to be centered around zero.

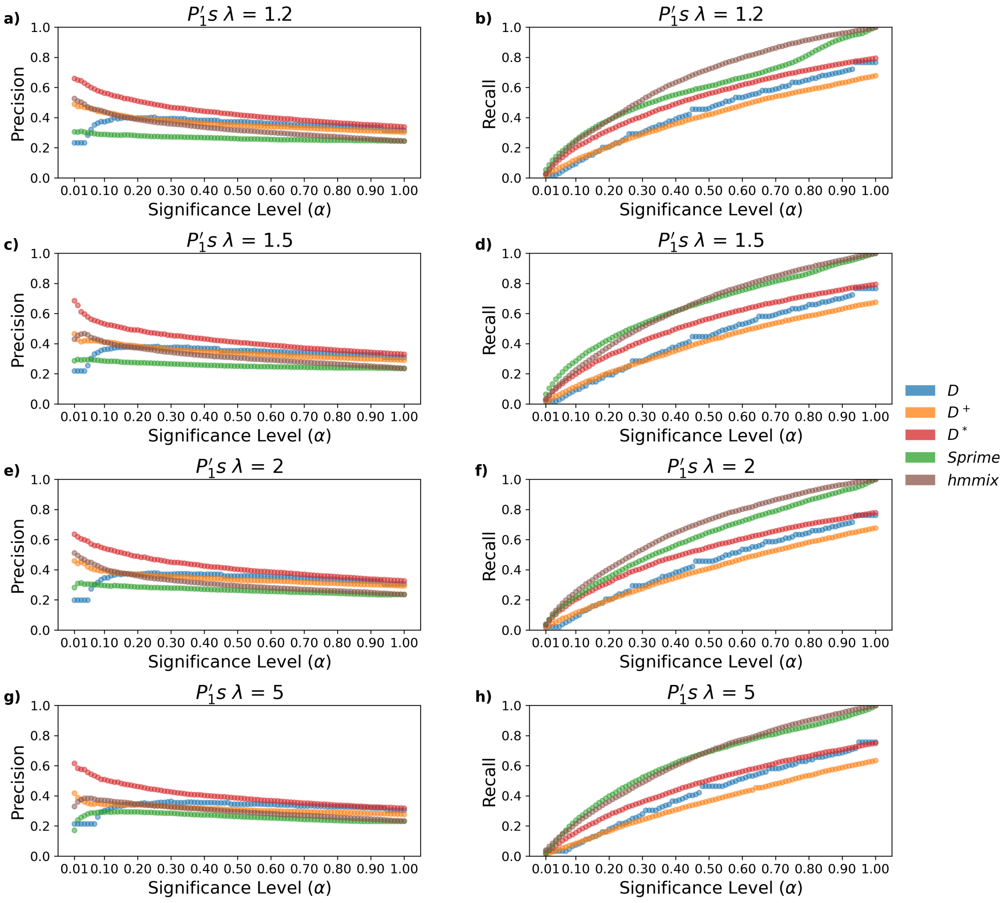


Figure S17: **Precision and Recall for Varying the Molecular Clock in** $P_1$ The precision and recall for identifying local introgression when varying the mutation rate by a factor of $\lambda$ in $P_1$. **a)** Precision and **b)** recall when $\lambda = 1.2$. **c)** Precision and **d)** recall when $\lambda = 1.5$. **e)** Precision and **f)** recall when $\lambda = 2$. **g)** Precision and **h)** recall when $\lambda = 5$.

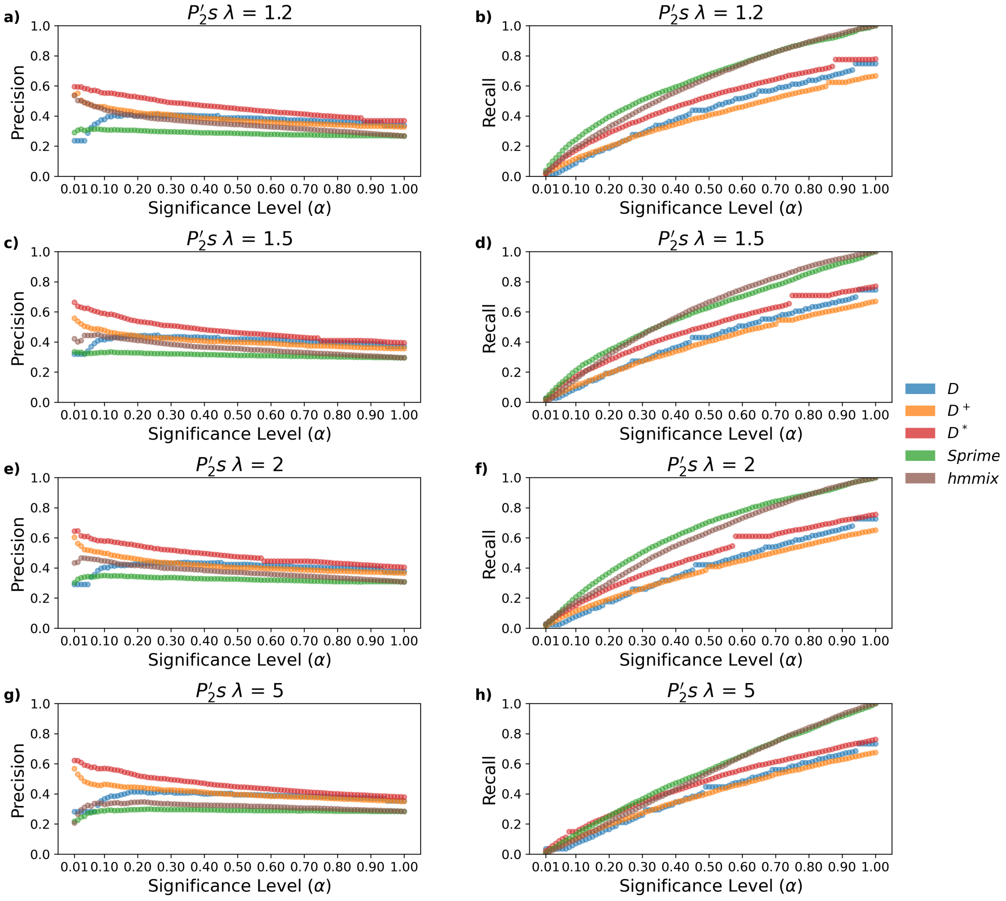


Figure S18: **Precision and Recall for Varying the Molecular Clock in $P_2$** The precision and recall for identifying local introgression when varying the mutation rate by a factor of $\lambda$ in $P_2$. **a)** Precision and **b)** recall when $\lambda = 1.2$. **c)** Precision and **d)** recall when $\lambda = 1.5$. **e)** Precision and **f)** recall when $\lambda = 2$. **g)** Precision and **h)** recall when $\lambda = 5$.

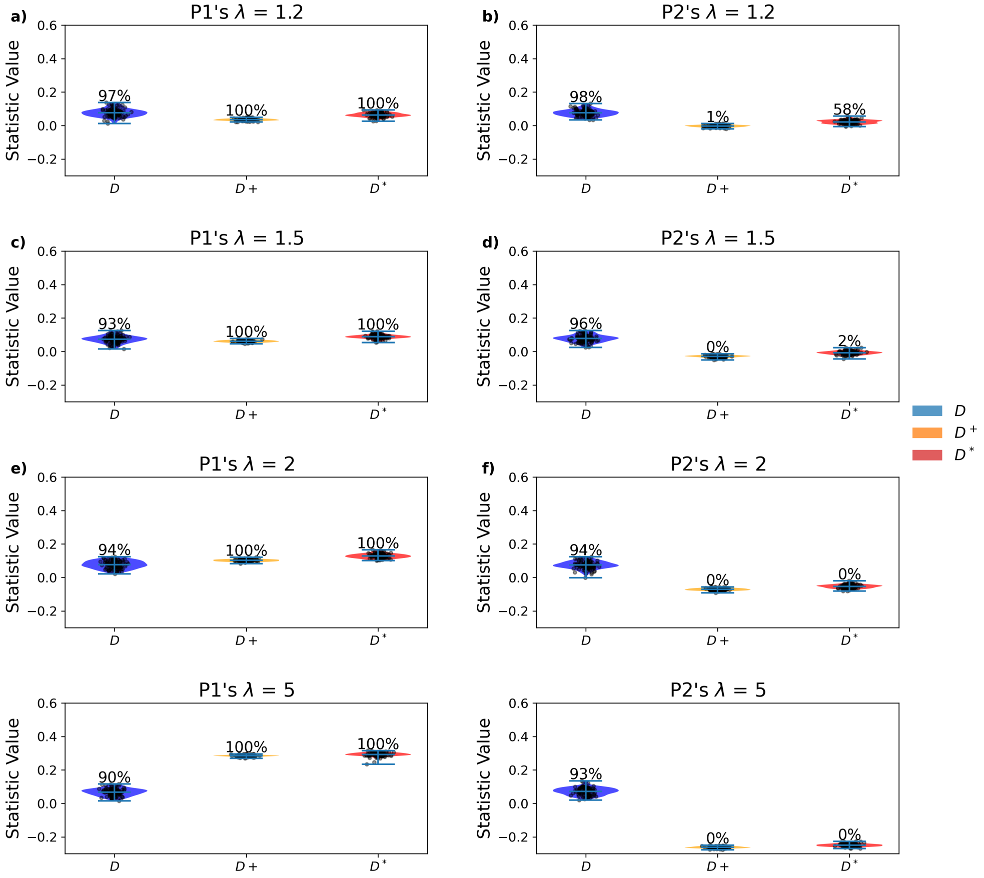


Figure S19: **Genome-Wide Power for Varying the Molecular Clock** The genome-wide power for introgression detection when the mutation rate in $P_1$ and $P_2$ by a factor of $\lambda$. Power for **a)** $P_1's\ \lambda = 1.2$, **b)** $P_1's\ \lambda = 1.5$, **c)** $P_1's\ \lambda = 2$, **d)** $P_1's\ \lambda = 5$, **e)** $P_2's\ \lambda = 1.2$, **f)** $P_2's\ \lambda = 1.5$, **g)** $P_2's\ \lambda = 2$, and **h)** $P_2's\ \lambda = 5$.

| | $N = 2500$ | $N = 5000$ | $N = 25000$ |
|---|---|---|---|
| P1 | $D$: 0.001/0.479<br>$D^{+}$: −0.002/0.147<br>$D^{*}$: 0.173/0.189 | $D$: 0.005/0.459<br>$D^{+}$: −0.000/0.142<br>$D^{*}$: 0.076/0.205 | $D$: 0.002/0.433<br>$D^{+}$: 0.001/0.135<br>$D^{*}$: −0.058/0.207 |
| P2 | $D$: −0.003/0.481<br>$D^{+}$: 0.001/0.147<br>$D^{*}$: −0.175/0.190 | $D$: 0.000/0.459<br>$D^{+}$: 0.001/0.142<br>$D^{*}$: −0.076/0.203 | $D$: 0.003/0.434<br>$D^{+}$: 0.000/0.135<br>$D^{*}$: 0.059/0.206 |

Table S4: **Mean and Standard Deviation of Null-Distribution for Varying Drift** The mean and standard deviation of the null distribution for $D$, $D^{+}$, and $D^{*}$ with varying drift in either population by changing the effective population size $N_e = \{2500, 5000, 25000\}$. We do not show the mean and standard deviation for *Sprime* and *hmmix* since they are not expected to be centered around zero.

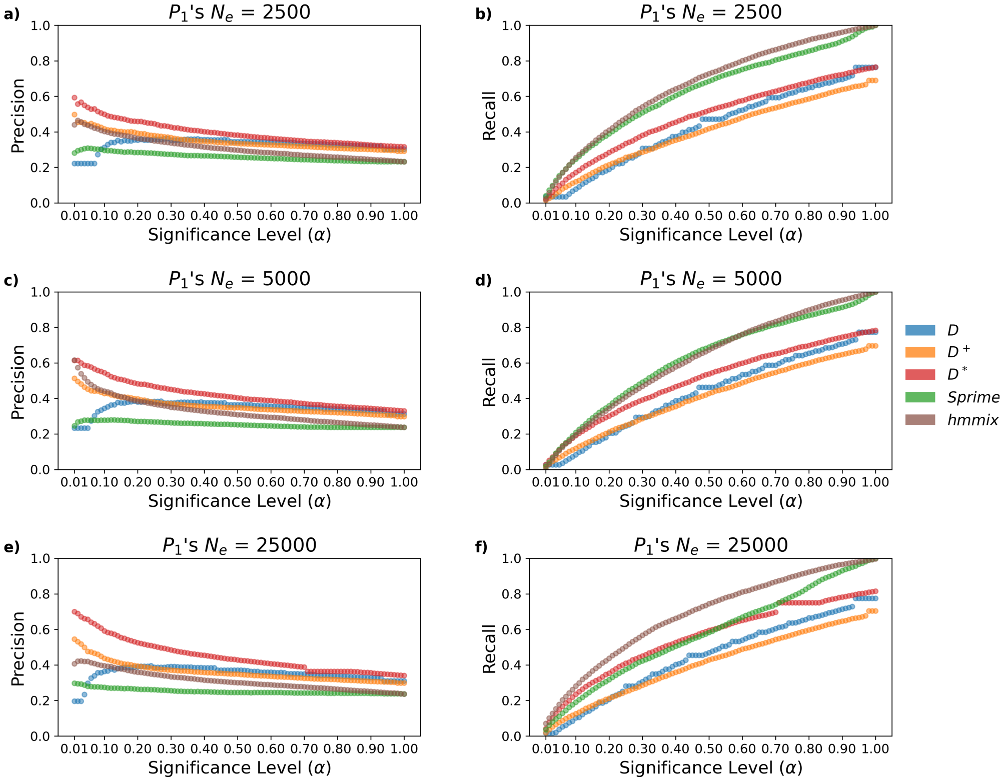


Figure S20: **Precision and Recall for Varying Drift in** $P_1$ The precision and recall for identifying local introgression with varying amounts of drift in $P_1$. **a)** Precision and **b)** recall when $P_1's$ $N_e = 2500$. **c)** Precision and **d)** recall when $P_1's$ $N_e = 5000$. **e)** Precision and **f)** recall when $P_1's$ $N_e = 25000$.

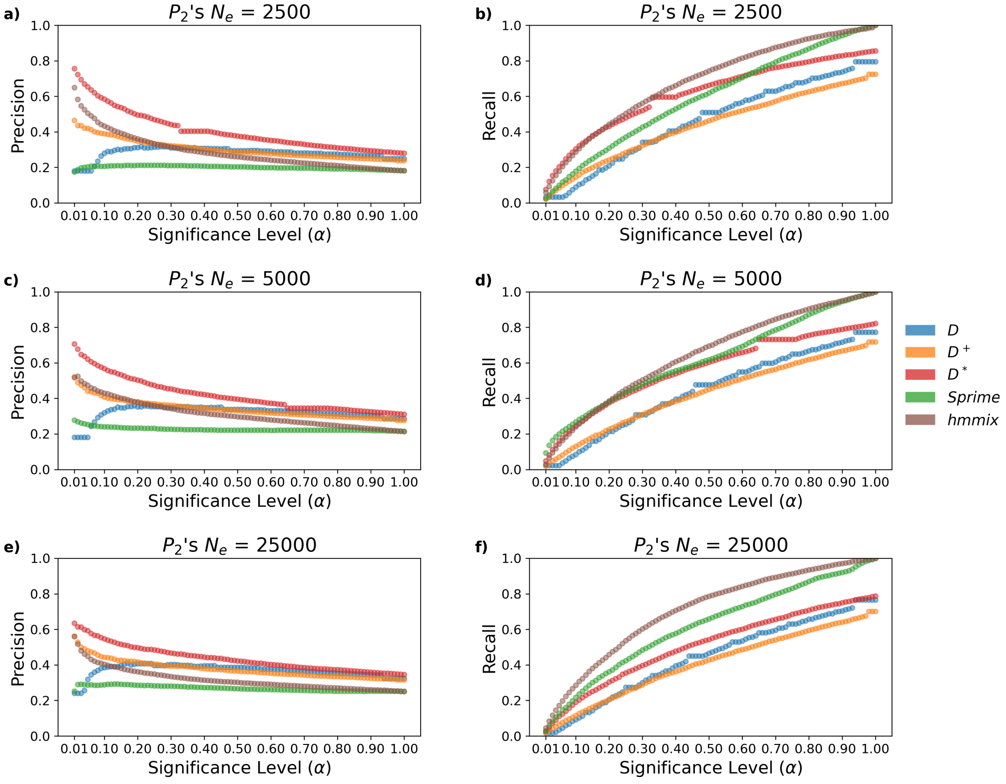


Figure S21: **Precision and Recall for Varying Drift in** $P_2$ The precision and recall for identifying local introgression with varying amounts of drift in $P_2$. **a)** Precision and **b)** recall when $P_2's$ $N_e = 2500$. **c)** Precision and **d)** recall when $P_2's$ $N_e = 5000$. **e)** Precision and **f)** recall when $P_2's$ $N_e = 25000$.

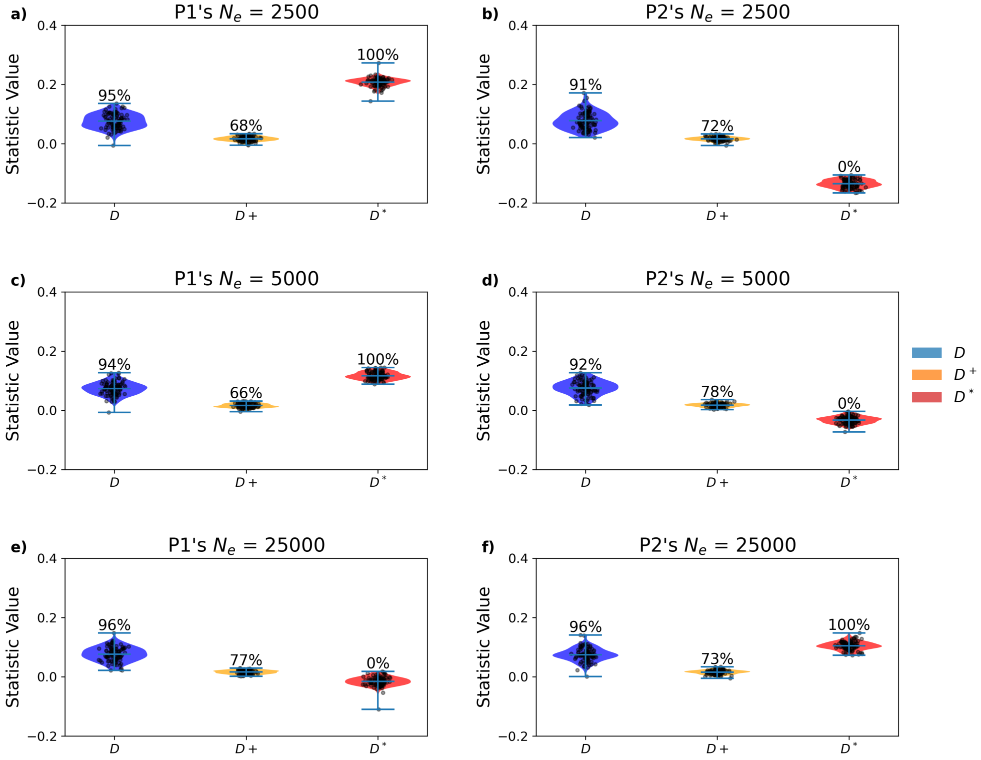


Figure S22: **Genome-Wide Power for Varying Drift** The genome-wide power for introgression detection when varying drift in $P_1$ and $P_2$. Power for **a)** $P_1's$ $N_e = 2500$, **b)** $P_1's$ $N_e = 5000$, **c)** $P_1's$ $N_e = 25000$, **d)** $P_2's$ $N_e = 2500$, **e)** $P_2's$ $N_e = 5000$, and **f)** $P_2's$ $N_e = 25000$.

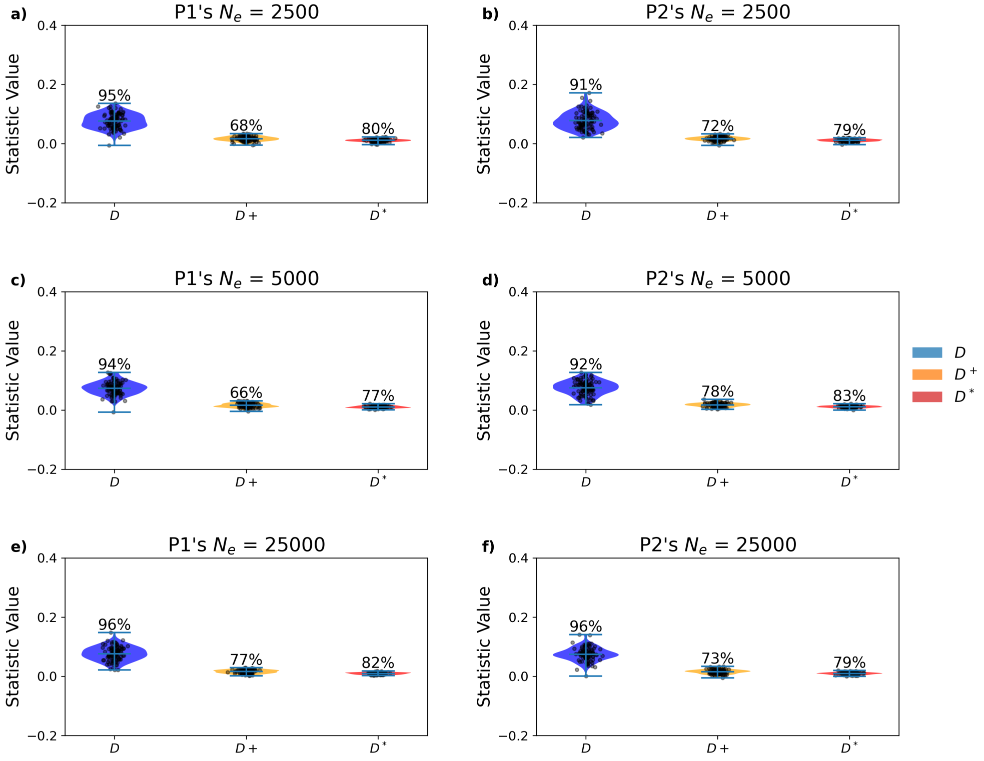


Figure S23: **Genome-Wide Power for Varying Drift with** $g = 1$ The genome-wide power for introgression detection when varying drift in $P_1$ and $P_2$ and the standardized sample size $g = 1$. Power for **a)** $P_1's\ N_e = 2500$, **b)** $P_1's\ N_e = 5000$, **c)** $P_1's\ N_e = 25000$, **d)** $P_2's\ N_e = 2500$, **e)** $P_2's\ N_e = 5000$, and **f)** $P_2's\ N_e = 25000$.

| | $N = 2500$ | $N = 5000$ | $N = 25000$ |
|---|---|---|---|
| P1 | D: $-0.000/0.021$<br>$D^{+}$: $-0.001/0.007$<br>$D^{*}$: $-0.000/0.004$ | D: $0.004/0.020$<br>$D^{+}$: $-0.000/0.006$<br>$D^{*}$: $0.000/0.004$ | D: $0.003/0.022$<br>$D^{+}$: $0.000/0.007$<br>$D^{*}$: $0.000/0.004$ |
| P2 | D: $-0.003/0.021$<br>$D^{+}$: $-0.001/0.007$<br>$D^{*}$: $-0.000/0.004$ | D: $-0.000/0.022$<br>$D^{+}$: $0.000/0.007$<br>$D^{*}$: $-0.000/0.004$ | D: $0.002/0.023$<br>$D^{+}$: $0.000/0.007$<br>$D^{*}$: $0.000/0.004$ |

Table S5: **Genome-Wide Mean and Standard Deviation for Varying Drift with $g = 1$ and No Introgression** The genome-wide mean and standard deviation without introgression when varying drift in $P_1$ and $P_2$ and the standardized sample size $g = 1$. Power for **a)** $P_1$'s $N_e = 2500$, **b)** $P_1$'s $N_e = 5000$, **c)** $P_1$'s $N_e = 25000$, **d)** $P_2$'s $N_e = 2500$, **e)** $P_2$'s $N_e = 5000$, and **f)** $P_2$'s $N_e = 25000$.

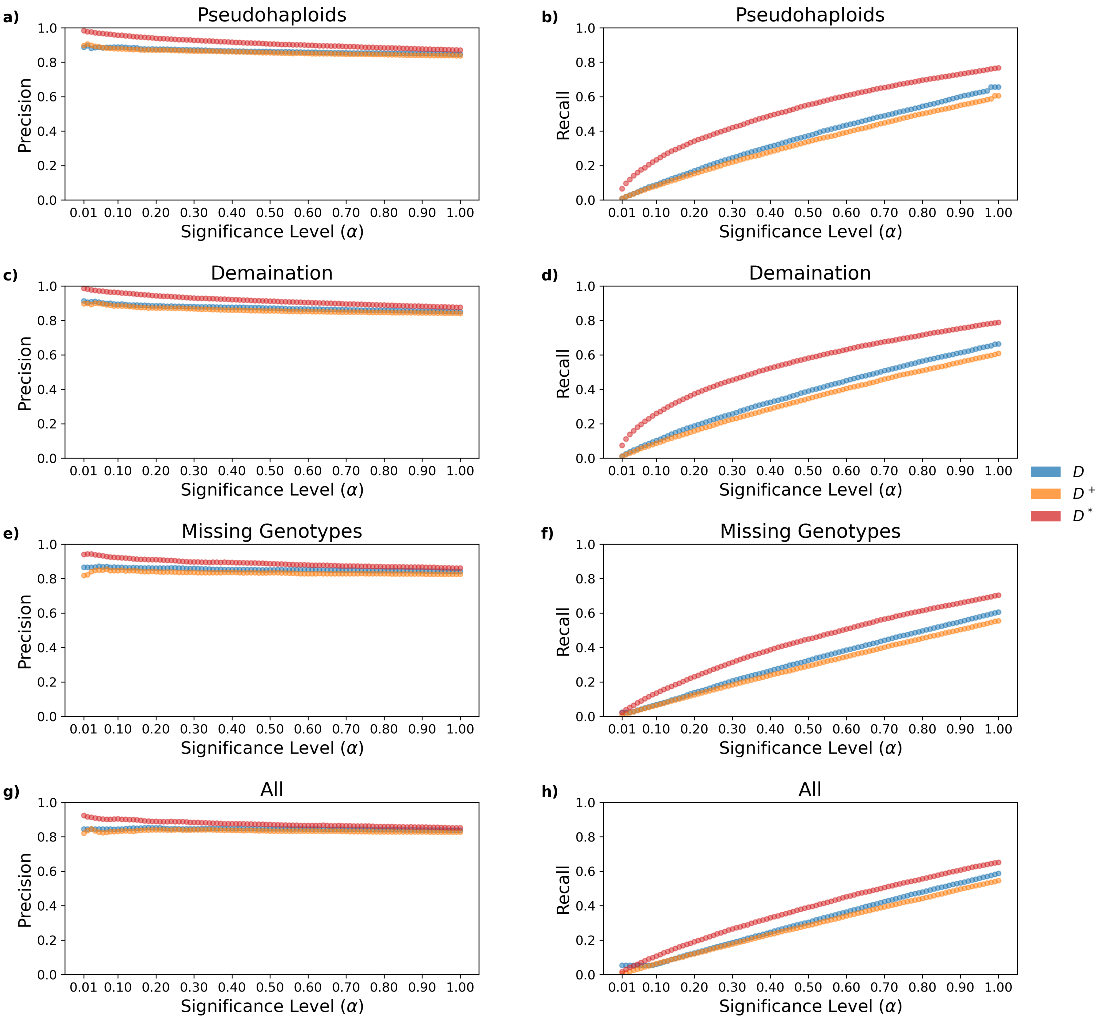


Figure S24: **Precision and Recall for Sources of Error in Ancient DNA** The precision and recall for identifying local introgression when sources of error common to ancient DNA are introduced. **a)** Precision and **b)** recall when pseudohaploids are introduced. **c)** Precision and **d)** recall when deamination is introduced. **e)** Precision and **f)** recall when missing genotypes are introduced. **g)** Precision and **h)** recall when pseudohaploids, deamination, and missing genotypes are introduced.

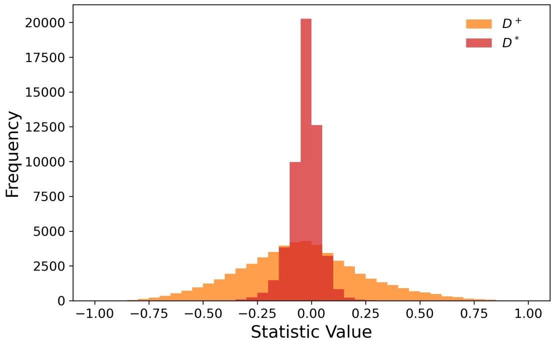


Figure S25: **Null Distribution for Denisovan and Papuan Introgression** The null-distribution for $D^+$ and $D^*$ in 50 Kb blocks using all Sardinian and Papuan samples. The mean and standard deviation for $D^+$ are -0.050 and 0.284, respectively. The mean and standard deviation for $D^*$ are -0.028 and 0.067, respectively.

Table S6: Top 0.1% of blocks with at least 10 sites and their overlapping protein coding genes exclusive to $D^*$ values on all Sardinian and Papuan samples.

| Region | $D^*$ | Genes |
|---|---|---|
| chr1:100500127-100549923 | 0.28183 | CDC14A,GPR88 |
| chr2:15350020-15399910 | 0.402892 | NBAS,RPS26P18 |
| chr2:15400032-15449785 | 0.34397 | NBAS |
| chr2:15450016-15499925 | 0.391287 | NBAS |
| chr2:131300151-131349960 | 0.294214 | PLEKHB2,FAR2P4,KLF2P4 |
| chr2:151700003-151749947 | 0.277121 | NEB |
| chr2:224000182-224049908 | 0.257698 | SERPINE2 |
| chr4:16750069-16799774 | 0.29699 | LDB2 |
| chr4:101400095-101449946 | 0.258302 | BANK1 |
| chr4:107700145-107749836 | 0.290816 | PAPSS1 |
| chr4:123250242-123299824 | 0.258087 | SPATA5 |
| chr4:123300742-123349869 | 0.302236 | SPATA5 |
| chr4:150700338-150749602 | 0.275387 | LRBA |
| chr4:151050038-151099860 | 0.251049 | RPS3A |
| chr5:54800401-54849824 | 0.360699 | CSPG4BP |

Table S6 – continued

| Region | $D^*$ | Genes |
|---|---|---|
| chr6:159850073-159899838 | 0.285552 | MAS1 |
| chr7:96300201-96349542 | 0.354934 | SLC25A13 |
| chr9:42900101-42949610 | 0.296936 | SNX18P5 |
| chr10:19350216-19399928 | 0.257331 | MALRD1 |
| chr10:73700011-73749562 | 0.291484 | SEC24C |
| chr10:74350281-74399904 | 0.256006 | ADK |
| chr10:74400047-74449866 | 0.273941 | ADK |
| chr10:74450171-74499778 | 0.284567 | ADK |
| chr11:4100485-4149958 | 0.247682 | RRM1 |
| chr11:78500470-78549716 | 0.290268 | NARS2 |
| chr12:10050040-10099910 | 0.257952 | CLEC9A,CLEC1A |
| chr12:20850050-20899851 | 0.346022 | SLCO1B3 |
| chr12:87900259-87949986 | 0.257881 | RPS4XP15 |
| chr14:23450291-23499621 | 0.265123 | NGDN |
| chr14:58250206-58299899 | 0.29471 | ARMH4,PSMA3,ARID4A |
| chr14:60950594-60999149 | 0.253788 | MNAT1,TRMT5,SLC38A6 |
| chr15:45250073-45299715 | 0.245841 | SLC28A2 |
| chr20:37200293-37249953 | 0.295835 | RPN2 |
| chr22:21450168-21496820 | 0.266034 | HIC2,TMEM191C |
| chr22:21500970-21547793 | 0.340411 | RIMBP3C |

Table S7: Top 0.1% of blocks with at least 10 sites and their overlapping protein coding genes according to $D^+$ values on all Sardinian and Papuan samples.

| Region | $D^+$ | Genes |
|---|---|---|
| chr2:15500113-15549983 | 0.827326 | NBAS |
| chr2:151750166-151800000 | 0.861246 | ARL5A |
| chr2:213600128-213649988 | 0.8334 | SPAG16 |
| chr3:51000035-51049848 | 0.853344 | DOCK3 |
| chr3:51050245-51099743 | 0.826569 | DOCK3 |
| chr3:69800121-69849828 | 0.8333 | MITF |
| chr3:71400504-71449831 | 0.839335 | FOXP1 |
| chr3:87750001-87799174 | 0.848428 | HTR1F |
| chr3:102250073-102299796 | 0.900797 | ZPLD1 |

Table S7 – continued

| Region | $D^+$ | Genes |
|---|---|---|
| chr3:102300814-102349721 | 0.851154 | ZPLD1 |
| chr4:94350137-94399948 | 0.883656 | RPL35AP11 |
| chr4:101200011-101249635 | 0.84719 | PPP3CA,MIR8066 |
| chr4:102750310-102799902 | 0.839853 | MANBA,UBE2D3 |
| chr5:65150389-65199914 | 0.823811 | ADAMTS6 |
| chr5:73700246-73749968 | 0.828556 | ARHGEF28 |
| chr6:56800008-56849290 | 0.839196 | DST |
| chr7:142750008-142799974 | 0.973808 | PRSS1,PRSS2,TRBD1,TRBJ1 |
| chr8:34950009-34999958 | 0.856505 | None |
| chr8:70700069-70749881 | 0.902988 | XKR9 |
| chr8:70750023-70799872 | 0.918332 | XKR9 |
| chr9:102600041-102649959 | 0.870202 | ZYG11AP1 |
| chr10:54250033-54299565 | 0.854832 | PCDH15 |
| chr11:26200323-26249614 | 0.826679 | ANO3 |
| chr11:32600161-32649567 | 0.855062 | EIF3M,CCDC73 |
| chr11:90650362-90699760 | 0.825971 | DISC1FP1 |
| chr12:93450066-93499734 | 0.827079 | MRPL42 |
| chr13:45400020-45449925 | 0.842676 | SLC25A30,PPIAP25 |
| chr16:59950170-59999894 | 0.835115 | LINC02141 |
| chr17:45600013-45649845 | 0.828391 | MAPK8IP1P2,RPS26P8 |
| chr17:45850342-45899977 | 0.841297 | MAPT-AS1,MAPT |
| chr18:46050146-46099729 | 0.832914 | PSTPIP2,RNY4P37,ATP5F1A |
| chr19:32600138-32649971 | 0.851545 | ANKRD27,SNORA68B,RPS12P31 |

Table S8: Top 0.1% of blocks with at least 10 sites and their overlapping protein coding genes according to *hmmix* values on all Sardinian and Papuan samples.

| Region | *hmmix* | Genes |
|---|---|---|
| chr2:176600001-176650000 | 0.7167965882352941 | MIR1246 |
| chr2:241750001-241800000 | 0.7135354117647059 | D2HGDH,GAL3ST2 |
| chr3:7650001-7700000 | 0.7121034117647058 | GRM7 |
| chr3:13350001-13400000 | 0.8121708235294117 | NUP210 |
| chr3:13400001-13450000 | 0.7719502352941177 | NUP210 |
| chr3:15650001-15700000 | 0.7603590588235294 | BTD,ANKRD28,MIR3134 |

Table S8 – continued

| Region | *hmmix* | Genes |
|---|---|---|
| chr3:15700001-15750000 | 0.8014722352941177 | BTD,ANKRD28 |
| chr3:15750001-15800000 | 0.7673574117647058 | ANKRD28 |
| chr3:15800001-15850000 | 0.7327869411764706 | ANKRD28 |
| chr3:15850001-15900000 | 0.7235469411764706 | ANKRD28,MIR563 |
| chr3:32050001-32100000 | 0.7782947058823529 | OSBPL10,NIFKP7 |
| chr3:58750001-58800000 | 0.7644494117647058 | CFAP20DC |
| chr3:58800001-58850000 | 0.7641554117647059 | CFAP20DC |
| chr3:76500001-76550000 | 0.7446471764705883 | ROBO2 |
| chr3:76550001-76600000 | 0.8099285882352941 | ROBO2 |
| chr3:151850001-151900000 | 0.7549878823529412 | SUCNR1 |
| chr3:175400001-175450000 | 0.74469 | NAALADL2 |
| chr3:175450001-175500000 | 0.7863381176470589 | NAALADL2 |
| chr4:2700001-2750000 | 0.8059950588235294 | FAM193A |
| chr5:113550001-113600000 | 0.7409244705882353 | YTHDC2 |
| chr5:150000001-150050000 | 0.7486089411764706 | TIGD6,HMGXB3 |
| chr5:150050001-150100000 | 0.7283549411764706 | HMGXB3,CSF1R |
| chr6:133850001-133900000 | 0.7788009411764706 | TARID,TCF21 |
| chr6:136000001-136050000 | 0.7233301176470588 | PDE7B |
| chr6:136150001-136200000 | 0.7960970588235294 | PDE7B |
| chr6:136200001-136250000 | 0.7229305882352941 | MTFR2 |
| chr6:136700001-136750000 | 0.7383697647058823 | MAP3K5 |
| chr7:64500001-64550000 | 0.7617342352941177 | ZNF680 |
| chr8:23100001-23150000 | 0.7559909411764706 | TNFRSF10C,TNFRSF10D |
| chr9:5950001-6000000 | 0.7614291764705882 | KIAA2026 |
| chr9:6000001-6050000 | 0.7094943529411765 | KIAA2026,MIR4665 |
| chr9:28100001-28150000 | 0.7314969411764707 | LINGO2 |
| chr9:28150001-28200000 | 0.7182455294117647 | LINGO2 |
| chr9:95400001-95450000 | 0.7467076470588235 | FANCC,PTCH1 |
| chr10:76550001-76600000 | 0.7137670588235294 | LRMDA |
| chr12:46750001-46800000 | 0.7164682352941176 | SLC38A4 |
| chr12:46800001-46850000 | 0.7770861176470588 | SLC38A4 |
| chr15:51850001-51900000 | 0.7078735294117647 | TMOD3 |
| chr15:54350001-54400000 | 0.7078501176470589 | UNC13C |
| chr15:58250001-58300000 | 0.7894623529411764 | ALDH1A2 |
| chr17:39600001-39650000 | 0.7358474117647059 | NEUROD2,PPP1R1B,STARD3 |
| chr18:63900001-63950000 | 0.7885099999999999 | SERPINB2,SERPINB10,HMSD |

Table S8 – continued

| Region | *hmmix* | Genes |
|---|---|---|
| chr21:14150001-14200000 | 0.7719110588235294 | LIPI |
| chr21:38650001-38700000 | 0.7358968235294118 | ERG |
